# SCG: Identifying spatially co-expressed genes using spatial covariance regression

**DOI:** 10.64898/2026.09.01.748618

**Authors:** Ihsan E. Buker, Yang Ni, Stephanie Hicks, Jian Kang, Satwik Acharyya

## Abstract

Spatial transcriptomics has enabled the advancement of gene expression analysis, yet spatial co-expression remains understudied. We introduce spatial covariance regression (SCR), a scalable Bayesian factor-model-based framework for estimation of spatially-resolved gene co-expression networks across tissue domains. These networks provide the spatial map of gene-gene correlations and enable the identification of spatially co-expressed genes (SCGs), which serve as potential prognostic biomarkers and therapeutic targets.

## Introduction

Spatially-resolved transcriptomics (SRT) technologies enable the measurement of gene expression while preserving the geospatial structure of the tissue microenvironment [1]. Disease progression is a result of the coordinated activity of a collection of genes, rather than genes in isolation. Network-based models, where genes are denoted as vertices and edges represent co-expression relationships, provide a natural framework for capturing the complex regulatory architecture of the transcriptome [2]. Several methods [2, 3] to recover gene co-expression networks estimate a single correlation network shared across the tissue domains, implicitly assuming a spatially homogeneous structure. The assumption of spatial homogeneity is often violated in tissues where co-expression varies across anatomical and pathological regions [4].

Recent efforts to account for the spatial heterogeneity of the tissue broadly fall into two classes: (i) study of marginal dependence via covariance matrices, and (ii) study of conditional dependence via precision matrices. Methods [4–6] to study marginal dependency are restricted to sample or spatial region level network inference and unable to provide spatially varying networks at the cellular resolution. Alternatively, graphical models [7, 8] are used to estimate the precision matrices, which characterize conditional dependence across the spatial domain of the tissue. These graphical models perform well for edge selection but struggle with edge estimation. Furthermore, they often require extensive post-processing to ensure the resulting precision matrices are symmetric and positive definite, neither of which is guaranteed by the modeling procedure. Jointly, no existing method delivers scalable, spot-resolution estimation of spatially varying marginal co-expression networks with rigorous uncertainty quantification.

To address these limitations, we propose spatial covariance regression (SCR) for estimation of spot-specific, spatially varying correlation matrices. Suppose a spatial tissue domain contains 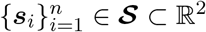 spatial locations, and we have the normalized gene expression profiles for *p* genes at spatial location ***s***_i_. Our goal is to estimate spatially varying *p*-dimensional networks where the marginal dependencies between genes are captured by an undirected graph G(***S***) = (V, E(***S***)) with a set of vertices V = {1, …, *p*} and a set of spatially varying edges E(***S***) ⊆ V × V. The spatial edge ***e***_gg*′*_ (***S***) ∈ E(***S***) connecting nodes *g* and *g*^*′*^ encodes their co-expression level across the spatial domain ***S***. To estimate E(***S***), we leverage a Bayesian spatial factor model to obtain a computationally tractable and low-dimensional structure. For further computational scalability, we decompose the loadings of the spatial factor model into two components: a spatial component (random-effects) that captures local variation through a set of basis functions, and a global component (fixed-effects) that regulates the connection of genes with basis elements along with factor scores, and a residual noise term (**Figure 1A**). We propose a mean-field variational approximation and develop a GPU-accelerated coordinate-ascent variational inference (CAVI) algorithm for scalability. SCR estimates spot-specific correlation matrices at each location, enabling the construction of spatially varying co-expression networks.

**Figure 1.**
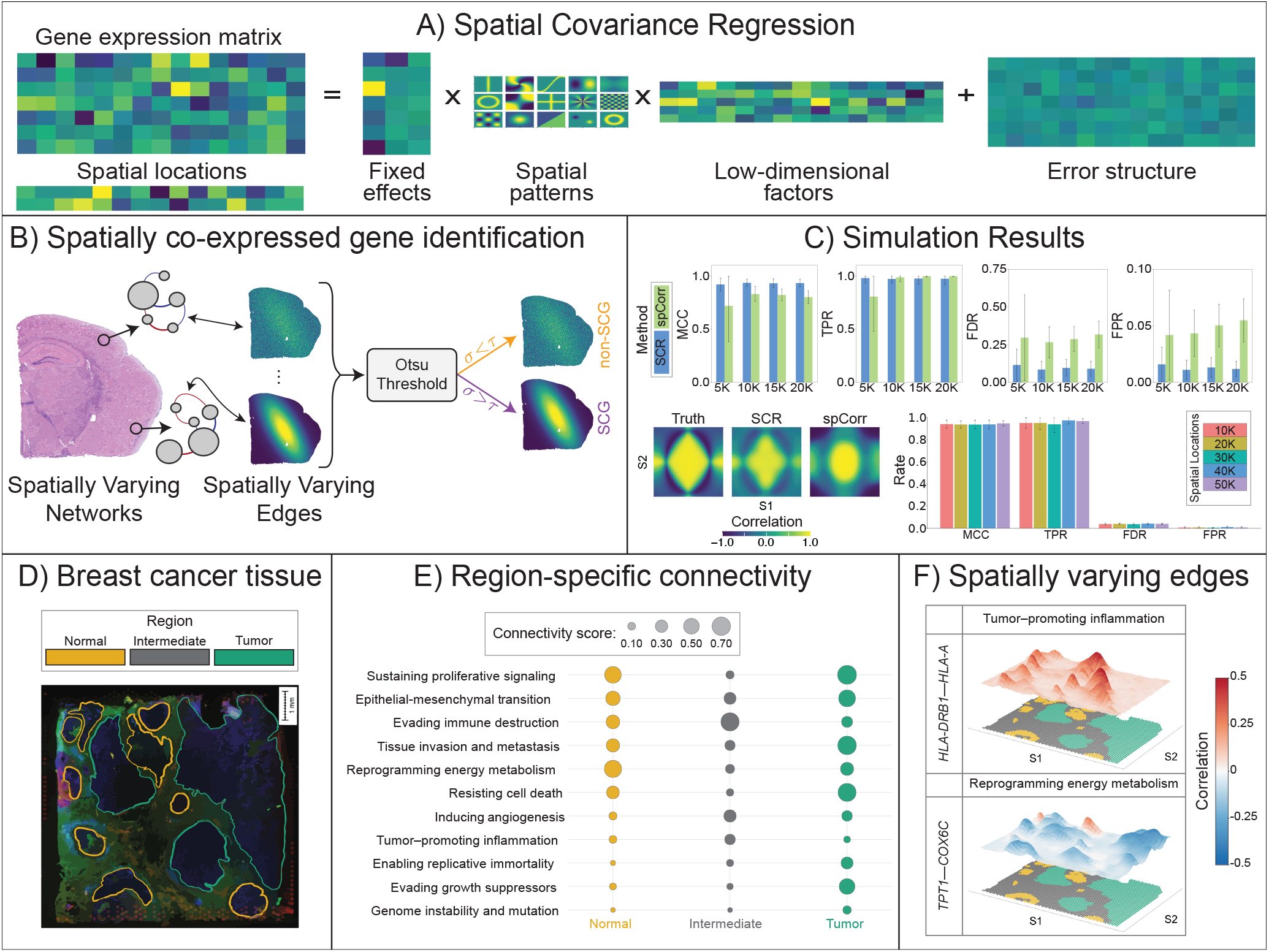
Spatial Covariance Regression (SCR) models spatially varying gene co-expression across tissue. (**A**), SCR decomposes the gene expression matrix into fixed effects, spatial random effects, low-dimensional factors, and residual error, enabling computationally-efficient, location-specific covariance estimation that allows for information sharing among spots and genes. (**B**), Spatially co-expressed gene (SCG) pairs are identified by applying Otsu thresholding to the posterior mean of the spatial standard deviation, yielding a threshold *τ*. Thereafter, the proportion of posterior samples in which the spatial standard deviation of an edge does not exceed *τ* is computed, yielding a posterior error probability (PEP). SCGs are then selected by Bayesian false discovery rate control at *α* = 0.1. (**C**), Benchmarking across simulated datasets of increasing size (5K–20K spatial locations) shows SCR outperforms spCorr across MCC, FDR, and FPR, with SCR recovering spatially varying correlation fields more faithfully than competing methods. With the exception of *n* = 5,000, spCorr attains marginally higher TPR compared to SCR; however, SCR considerably outperforms spCorr when considering balanced classification, as indicated by superior MCC across sample sizes. We further examine the performance of SCR for higher-resolution datasets (10K–50K spatial locations) to which spCorr failed to scale. We see strong, consistent performance by SCR across sample sizes (See Supplementary Section S2 for further details). (**D**), Application to a human breast cancer tissue section with three pathologist-annotated regions (Normal, Intermediate, tumor). (**E**), Region-specific gene network connectivity scores across Hallmarks of Cancer pathways reveal progressive rewiring from normal to tumor tissue, with angiogenesis and proliferative signalling most activated in the tumor region. (**F**), Spatially varying correlation fields for two gene pairs (*HLA-DRB1* –*HLA-A* and *TPI1* –*COX6C*) involved in tumor-promoting inflammation and reprogramming energy metabolism, respectively.

Using the estimated networks, we define spatially co-expressed genes (SCGs) as gene pairs corresponding to edges in E(***S***) where the correlations exhibit significant spatial variation (**Figure 1B**). An Otsu threshold [9] is applied to the posterior mean spatial standard deviation of edges to separate edges with negligible versus substantial spatial variation. Thereafter, we compute the posterior probability that an edge does not exceed this threshold (posterior error probability, PEP). In this context, the PEP corresponds to the probability of erroneous classification of an edge as an SCG given the data. The set of SCGs is constructed using FDR control, which involves ranking the edges in ascending order of PEPs and selecting the largest number of edges *k*^⋆^ such that the mean PEP of the first *k*^⋆^ edges is less than or equal to a pre-specified FDR level *α* [10]. Further methodological details are provided in **Methods Section** and **Supplementary Section S1**.

## Results

Analogous to spatially variable genes (SVGs [11]), which capture the spatial variation in gene expression, SCGs capture the spatial variation in gene co-expression. Given the intrinsically coordinated nature of gene regulatory programs, co-expression analysis will enable a system-level understanding of signaling pathways and disease etiology. By estimating a correlation matrix at every location, SCR incorporates the spatial heterogeneity of gene co-expression networks across the tissue microenvironment [12]. SCG identification is then an inferential procedure applied to these estimated networks, in which an edge is classified as an SCG according to the FDR-based rule. To assess the performance of SCR, we performed extensive simulation studies comparing it against existing methods (spCorr [6]) with respect to SCG identification and correlation surface recovery (**Figure 1C, Supplementary Section S2**).

We applied SCR to SRT data from a HER2-amplified invasive ductal carcinoma sample [13] which is delineated with pathologist annotations as tumor, intermediate, and normal regions (**Figure 1D**). At each spot, SCR estimates a correlation matrix encoding network edges and enables the quantification of spatially varying connectivity scores (CS) to study pathway level activity across the tissue domain. The CS [7] is computed as the number of edges in a given network that exceed 0.1 in magnitude, normalized by the maximum count across spots and pathways to yield a score that ranges from 0 (low connectivity) to 1 (high connectivity). We computed CS for the ten cancer hallmark pathways [14] and the epithelial–mesenchymal transition program [15] (**Figure 1E, Table S10**).

SCR identified the sustaining proliferative signaling pathway with the strongest connectivity in the tumor region (CS: 0.75), which is a known driver of proliferative signaling for HER2 amplification [16]. The tissue invasion and metastasis pathway has a strong CS in the tumor region (CS: 0.73) and HER2 overexpression is known to promote breast cancer cell invasion and metastasis [17]. Both pathways have moderately low levels of connectivity in normal (CS 0.58, 0.36, respectively) and intermediate region (CS 0.12, 0.20, respectively). These trends are consistent with fundamental mechanisms of tumor biology that the coordinated activity of these pathways drive tumorigenesis, promoting key hallmarks of cancer [14].

We identified SCGs from each of the 11 pathways (**Supplementary Table S9**). Here, we focus on the *HLA-DRB1* –*HLA-A* edge from the tumor-promoting inflammation pathway and the *TPI1* –*COX6C* edge from the reprogramming energy metabolism pathway (**Figure 1F**). The *HLA-DRB1* – *HLA-A* edge was characterized by strong positive correlation localized along the tumor boundary, with practically no correlation within the tumor core. *HLA-DRB1* and *HLA-A* encode MHC class II and class I molecules, respectively, which present antigen to T cells and these classes of genes were found to co-occur in SRT [1]. The loss of co-expression in the tumor core is consistent with tumors evading immune destruction [18]. The *TPI1* –*COX6C* edge was characterized by broad negative correlation across the tissue, largest in magnitude within the tumor core. *TPI1* and *COX6C* encode a glycolytic enzyme and a Complex IV subunit of the oxidative-phosphorylation chain, respectively. Glycolytic and oxidative-phosphorylation programs are negatively correlated across tumor transcriptomes [19]. This trade-off is spatially organized with glycolysis favored in the hypoxic tumor core [20]. The trends identified by SCR suggest that glycolysis and oxidative-phosphorylation are inversely co-regulated where the metabolic trade-off is most active. Together, these edges demonstrate SCR resolving both a positive, immune-associated coupling localized along the tumor boundary and a negative, metabolic coupling localized to the tumor core, recovering established tumor immunobiology and metabolic reprogramming at spatial resolution (**Supplementary Figures S14–S16**).

Using SCR, we analyzed Alzheimer’s disease (AD) mouse-model brain data generated with the 10x Genomics Xenium platform, comparing a 13.4-month-old wild-type (WT) mouse with a 17.9-month-old TgCRND8 transgenic (MUT) mouse with advanced *β*-amyloid deposition [21]. For both samples, we focused on the plaque-induced gene (PIG) module [22], which is a co-expression program of genes induced near amyloid plaques. We restricted the analysis to the 46 PIG genes present in the Xenium panel across ~ 55,000 cells for both samples. We classified 301 and 249 SCGs in the MUT and WT samples, respectively. The higher number of SCGs identified in the transgenic brain with high *β*-amyloid load is consistent with the original finding that PIG co-expression intensifies with plaque burden [22].

Figure 2A shows three edges from the PIG module, each projected onto the spatial coordinates and anatomical annotation of the WT and MUT sections. Only *Ctsd* –*Ctsb* was identified as an SCG in both samples; the remaining two edges were identified as SCGs only in the MUT sample. The *Ctsd* –*Ctsb* edge was positively co-expressed in both genotypes but with distinct topology. In WT the positive co-expression was confined to the striatum, whereas in MUT it was broad and strong. The *C1qa*–*Apoe* and *Lyz2* –*Apoe* edges were positively co-expressed across most of the brain, with a localized region of negative co-expression in the lateral septal complex, hypothalamus, midbrain, and hindbrain. Further edges from both samples are provided in **Supplementary Figures S19, S20, S23, S24**.

**Figure 2.**
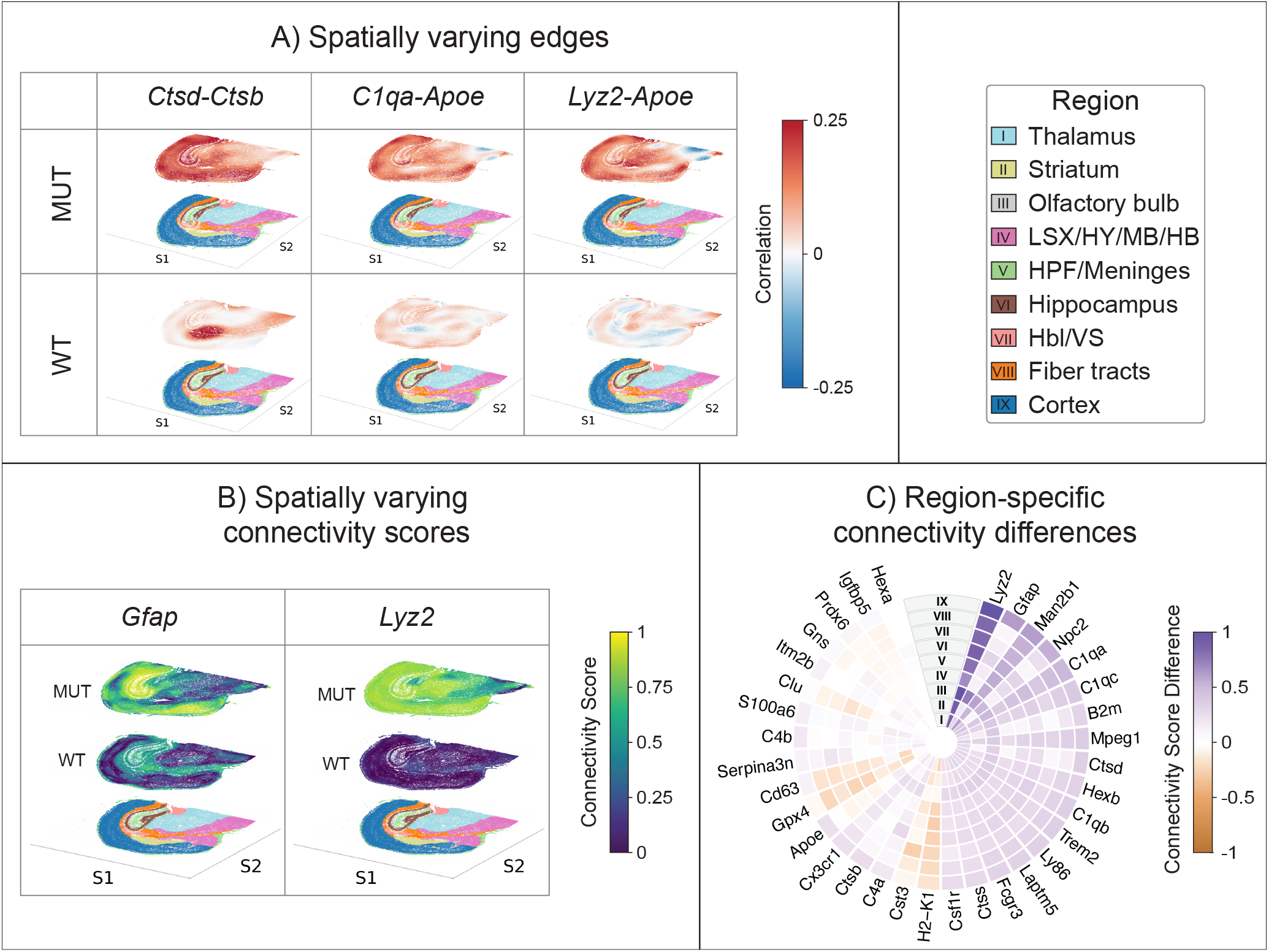
SCR reveals spatially varying gene co-expression in a mouse model of AD. (**A**), Spatially varying correlation fields for three gene pairs (*Ctsd* –*Ctsb, C1qa*–*Apoe*, and *Lyz2* –*Apoe*) compared between mutant (MUT) and wild-type (WT) mice. With the exception of *Ctsd* –*Ctsb*, for which the edge was spatially varying in both MUT and WT samples, the two remaining pairs were SCGs only for the MUT sample and not for the WT sample. Co-expression patterns are spatially structured and show a consistent relationship to underlying anatomical regions, with MUT typically exhibiting elevated positive correlations relative to WT. (**B**), Spatially varying connectivity scores for *Gfap* and *Lyz2* across the tissue, with MUT mice displaying broader and stronger connectivity relative to WT. (**C**), Region-specific connectivity differences between MUT and WT across nine anatomical regions (I–IX), computed for all genes in the PIG pathway that were involved in spatially varying edges for both samples. Positive scores (purple) indicate MUT-enriched connectivity; negative scores (brown) indicate WT-enriched connectivity. Complement genes (*C1qa, C1qb, C1qc, C4b*), lysosomal genes (*Ctsd, Hexb*), and disease-associated microglial markers (*Trem2, Apoe*) show notable upregulation in the MUT sample. The foregoing upregulation is most consistently noted in the cortical regions. Since the PIG pathway is upregulated near *β*-amyloid plaques, a plausible hypothesis is that the cortex suffers from heavier plaque burden.

For a given gene, we define its connectivity score (CS) at each spot based on the number of significant edges. Figure 2B displays the spatially resolved CS of *Gfap*, an astrocyte marker [23], and *Lyz2*, a marker of myeloid lineage cells [24]. In WT, *Gfap* connectivity was low in the thalamus and cortex but moderate in the remaining regions. Conversely, in MUT, *Gfap* connectivity increased across most regions. *Lyz2* connectivity, by contrast, was uniformly low across the brain in WT but broadly high in MUT. Aggregating connectivity differences by anatomical region (Figure 2C) shows that this reorganization is gene- and region-specific rather than uniform. Increases were most prominent in markers of activated myeloid cells [25] and the plaque-associated complement response [26], such as *Lyz2, Trem2*, and *C1qa*. The spatially varying connectivity scores for SCGs are shown in **Supplementary Figures S21–S22**.

## Discussion

Our proposed SCR framework provides rigorous estimation of spatially varying correlation surfaces and conceptualizes the novel idea of SCGs. Instead of the study of mean expression and identification of SVGs, we focus on spatially varying correlation structure to classify gene interactions as SCGs. This enables us to study pathway level signaling across the spatial regions of the tissue microenvironment. The GPU-accelerated CAVI implementation ensures scalability of the method with high-dimensional ST data.

Several extensions of SCR can be considered. A multi-sample formulation could infer consensus SCGs across individuals while accounting for inter-subject variability, enabling population-level co-expression studies and cohort comparisons relevant to disease stages and biomarker discovery. For further scalability, one can replace the CAVI algorithm with a stochastic variational inference algorithm, which will enable SCR to scale with ST datasets with millions of cells, as produced by the latest imaging-based platforms.

## Methods

### SCR model

Suppose a spatial tissue domain contains ***s***_i_ ∈ ***S*** ⊂ ℝ_2_ for *i* = 1, …, *n* spatial locations or cells, and let ***y***(***s***_i_) = (*y*_i1_, …, *y*_ip_)^*⊤*^ ∈ ℝ_p_ denote the normalized gene expression profiles for *p* genes at location ***s***_i_. We aim to estimate a spatially varying *p*-dimensional network for which the marginal dependencies between genes are captured by a spatially varying undirected graph G(***S***) = (V, E(***S***)) with vertex set V = {1, …, *p*} and a spatially varying edge set E(***S***) ⊆ V × V. An edge in E(***S***) encodes the coexpression level of two genes at location ***s***_i_, which we quantify through the correlation coefficient.

To estimate E(***S***), we assume ***y***(***s***_i_) ~ N_p_(0, **Σ**(***s***_i_)) for each location ***s***_i_, where **Σ**(***s***_i_) characterizes the marginal dependencies among genes and is consistent with the corresponding graph structure. To accommodate the sparse, high-dimensional nature of spatial genomics data, we use a spatial factor model

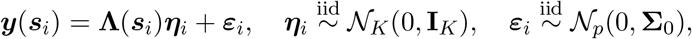

where **Λ**(***s***_i_) ∈ ℝ_p*×*K_ is the spatial factor loading matrix, ***η***_i_ ∈ ℝ_K_ are the latent factor scores, and ***ε***_i_ ∈ ℝ_p_ are zero-mean Gaussian errors with covariance **Σ**_0_ ∈ ℝ_p*×*p_. A low-dimensional structure is induced by the standard factor model assumption, i.e., *K* ≪ *p*.

Our goal is to estimate spatially varying gene coexpression networks. At each spatial location ***s***_i_, the edges of the undirected graph G(***S***) = (V, E(***S***)) are characterized by pairwise associations derived from **Σ**(***s***_i_). In particular, we quantify the spatially varying coexpression between genes *g* and *g*^*′*^ using the correlation coefficient,

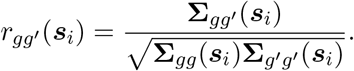

This yields a spatially varying correlation network that captures changes in gene coexpression across the tissue domain. Further details about model specifications regarding low-rank decomposition and identifiability are provided in **Supplementary Section S1.1**.

### Prior specification

To develop a parsimonious and scalable modeling framework, we decompose the spatial factor loading matrix into a global–local structure, representing the factor loadings as weighted combinations of a reduced set of spatial basis elements as

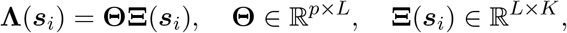

where **Θ** captures global gene–basis associations and **Ξ**(***s***_i_) is a collection of nonparametric spatial basis functions that encode local spatial variation [27]. This decomposition reduces the number of parameters required to model **Σ**(***s***_i_) for a spatial location *s*_i_ from *p*(*p* + 1)*/*2 for a fully parameterized covariance to *pL* global loading parameters, *LK* spatial basis functions, and *p* residual variances, reducing the parameter complexity from quadratic to linear in *p* when *K* ≪ *p* and *L* ≪ *p*. This dimension reduction technique enables the model to scale efficiently for high-dimensional spatially varying gene networks.

We place a multiplicative gamma process shrinkage prior [28] on **Θ** as

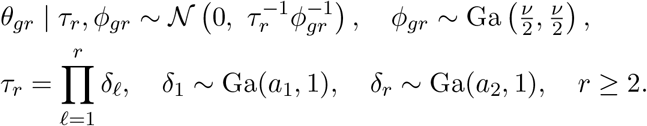

Here *ϕ*_gr_ is a local (gene–basis-specific) precision and *τ*_r_ is a global column precision. This prior is motivated by three considerations: (1) it imposes increasing shrinkage across the columns of **Θ**, allowing *L* to be chosen conservatively large while redundant spatial bases are effectively truncated; (2) it permits a small subset of genes to have significant loadings while strongly shrinking the remaining towards zero to mimic the sparse structure of SRT; and (3) it complements the loading decomposition by sharing information across genes and spatial locations while concentrating posterior mass on an effectively lower-dimensional structure.

For the spatial basis functions, we assume independent zero-mean Gaussian process (GP) priors to model the functions nonparametrically. For each basis–factor pair (*r, k*), we place the GP prior as

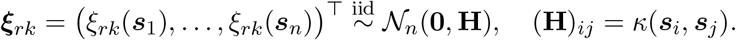

The kernel *κ* governs the smoothness and spatial range of the learned bases and plays a central role in capturing latent spatial structure. We use a rational quadratic (RQ) kernel,

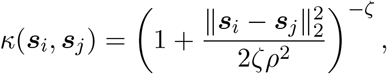

which allows correlations to decay at multiple spatial scales, accommodating both broad and localized expression patterns. The RQ kernel can be viewed as a scale mixture of squared-exponential (SE) kernels over a distribution of length-scales: *ρ* controls the rate at which correlation decays with 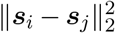, while *ζ* governs the implied length-scale mixture. Smaller *ζ* places more weight on shorter length-scales, permitting greater local variation while preserving long-range trends, and as *ζ* → ∞ the RQ kernel converges to the SE kernel [29].

To complete prior specification, we assume the residual covariance 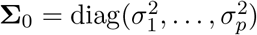 is diagonal and place independent conjugate priors on the gene-specific residual precisions,

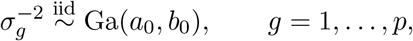

equivalently inverse-gamma priors on the variances 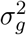. This captures gene-specific residual variability while retaining conjugacy for the coordinate-ascent updates.

### Variational inference

Bayesian inference for factor models is often performed using Markov chain Monte Carlo (MCMC), but such algorithms scale poorly in high-dimensional settings [30]. For computational efficiency, we develop a coordinate ascent variational inference (CAVI) algorithm for estimation and uncertainty quantification of the model parameters

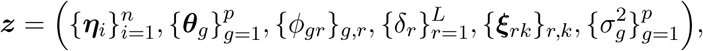

where ***θ***_g_ denotes row *g* of **Θ**. Variational inference posits a family of densities *Q* over ***z*** and seeks the surrogate posterior *q*(***z***) ∈ *Q* minimizing the Kullback–Leibler divergence to the true posterior,

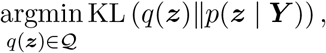

equivalently maximizing the evidence lower bound (ELBO),

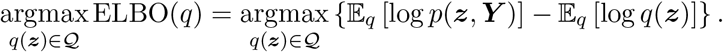

We adopt a mean-field variational family

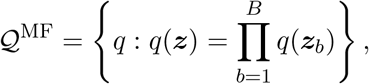

which partitions the parameters into *B* disjoint blocks [31]. Letting ***z***_*−*b_ denote all blocks except ***z***_b_, the optimal variational factor is

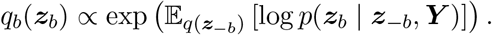

When the complete conditional belongs to the exponential family, the optimal factor remains in the same family, reducing ELBO maximization to updating the associated natural parameters. Complete derivations are provided in **Supplementary Section S1.2**.

### SCG identification

We estimate the spatially varying covariance matrices as

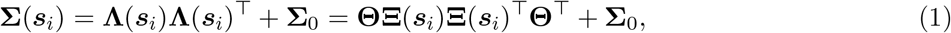

which quantify the marginal associations between genes at location ***s***_i_. To compute the posterior mean correlation fields, let

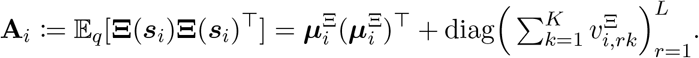

With ***µ***_Θ_ = E_q_[**Θ**] and 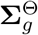 the variational covariance of row *g* of **Θ**, the posterior mean covariance is

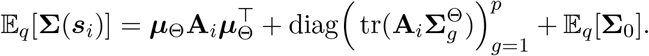

For each gene pair (*g, g*^*′*^), the posterior mean correlation at location ***s***_i_ is

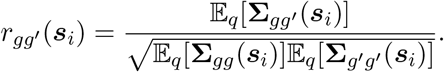

Let ***r***_gg*′*_ = *r*_gg*′*_ (***s***_1_), …, *r*_gg*′*_ (***s***_n_) _*⊤*_ denote the resulting length-*n* correlation field. To quantify spatial variability while accounting for posterior uncertainty, we approximate

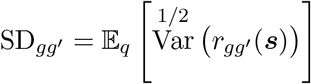

via Monte Carlo sampling from the variational posterior: we draw *M* independent samples 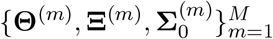 and form **Λ**^(m)^(***s***_i_) = **Θ**^(m)^**Ξ**^(m)^(***s***_i_). For each gene pair and draw *m*, we compute

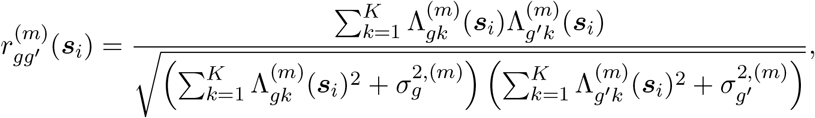

the spatial standard deviation 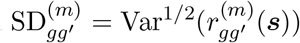, and the estimate 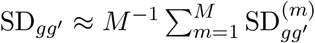. We apply Otsu’s thresholding method to {SD_gg*′*_}_g<g*′*_ to obtain a data-driven threshold *τ* [9], then compute

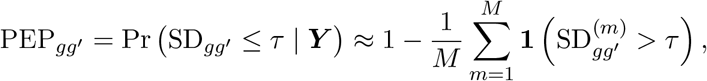

and label gene pairs as spatially co-expressed genes (SCGs) when PEP_gg*′*_ is sufficiently small, while controlling the false discovery rate.

### Simulation study

We conducted an extensive simulation study to assess the performance of SCR in graph structural recovery and edge estimation. We compared SCR against spCorr [6] across spatial grids of 5,000 to 50,000 locations with *p* = 30 genes and 20 independent replicates. Structural recovery was assessed using the Matthews correlation coefficient (MCC), true positive rate (TPR), and false positive rate (FPR). Edge estimation was assessed using the RV coefficient [31], which measures concordance between estimated and true spatial correlation matrices at each location. The RV coefficient takes values in [0, 1], with values near 1 indicating a high level of agreement. Data were generated from a spatial factor model with a certain number of spatially varying and spatially constant latent factors. Spatial factor fields were constructed by sampling from a library of 100 non-parametric functions.

mimicing the sparse nature of SRT data, we designed the gene loadings to be sparse and heterogeneous. A small fraction of genes loaded on spatial factors with coefficients from N (0, 1) and a fraction loaded on constant factors with coefficients from N (0, 0.01). All remaining genes had zero loadings. Following the equation in (1) of location-specific covariance, we generate the gene expression from ***y***(***s***_i_) ~ N_p_(**0, Σ**(***s***_i_)). Ground-truth SCG labels were assigned by applying Otsu’s threshold to the spatial standard deviation of the true correlation field for each gene pair. All the results are summarized across 20 replicated studies.

SCR was benchmarked against spCorr under the identical data-generating mechanism across the spatial resolutions for which spCorr was computationally feasible (*n* ≤ 20,000). spCorr did not scale beyond 20,000 locations, whereas SCR was scalable up to 50,000. At every matched sample size, SCR recovered the spatially varying network more precisely (Figure 1C, top row) with higher mean MCC (0.920, 0.935, 0.931, 0.933 at *n* = 5,000, 10,000, 15,000, 20,000) than spCorr (0.719, 0.830, 0.821, 0.801). spCorr achieved marginally higher TPR with over-selection of edges and at the cost of high FPR. SCR incurred only a marginal reduction in TPR while maintaining a substantially lower FPR. This trade-off was well captured by the MCC, which balances the two metrics. SCR also performed better in terms of edge estimation as shown in Figure 1C and **Supplementary Figures S1–S4**. The inferential objects of interest in spCorr are individual spatially varying edges, which are estimated independently and can be arbitrarily parallelized across CPUs. In contrast, SCR estimates an entire spatially varying network jointly. Because the two methods operate on different computational units, comparing their raw wall-clock times directly would be misleading. Therefore, we placed both methods on a common per-edge scale. For SCR, we recorded the total wall-clock time for each of the 20 replicates and averaged across them, yielding mean times ranging from approximately 9.6 minutes at the smallest grid to 291 minutes at the largest. We then divided each mean time by the 435 edges, corresponding to all pairwise edges among the *p* = 30 genes. This yielded the average per-edge run times. On this per-edge basis, SCR was substantially faster than spCorr (1.3, 2.8, 4.7, and 7.1 seconds for SCR at *n* = 5,000, 10,000, 15,000, and 20,000, versus 8.9, 19.4, 32.8, and 37.1 seconds for spCorr). Moreover, SCR’s run times were far more consistent, as shown by the considerably narrower error bars (**Supplementary Figure S13**). Full numerical results and simulation figures are provided in **Supplementary Section S2**.

For different numbers of spatial locations *n* ∈ {10,000, 20,000, 30,000, 40,000, 50,000}, SCR demonstrated consistently high performance in structural recovery and edge estimation. The mean MCC and TPR are {0.940, 0.936, 0.935, 0.937, 0.946} and {0.950, 0.950, 0.938, 0.973, 0.967} respectively while maintaining a low FPR {0.007, 0.008, 0.006, 0.011, 0.008} and FDR {0.039, 0.040, 0.038, 0.042, 0.041}. The mean RV coefficient ranged from 0.984 to 0.990 across different numbers of spatial locations. These results demonstrate that the proposed method consistently recovers the covariance structure across the spatial domains.

## Data availability

The breast cancer dataset was obtained from the BayesSpace manuscript [13]. The Alzheimer’s disease dataset was similarly obtained from the 10x Genomics Datasets [portal](https://www.10xgenomics.com/datasets/xenium-in-situ-analysis-of-alzheimers-disease-mouse-model-brain-coronal-sections-from-one-hemisphere-over-a-time-course-1-standard). For the TgCRND sample, we analyzed the 17.9-month time point, and for the wild-type sample, we analyzed the 13.4-month time point. Both datasets and supporting materials have been deposited in Zenodo and are publicly available under DOI: 10.5281/zenodo.22061520.

## Code Availability

SCG is available as an open-source Python package at https://github.com/iebuker/SCG. Documentation and tutorials are available at https://iebuker.github.io/SCG/.

