## Supplementary Material for "SCG: Identifying spatially co-expressed genes using spatial covariance regression"

### Supplementary for “SCG: Spatially Co-Expressed Gene Identification through Spatially Varying Networks”

### S1 Spatial Covariance Regression (SCR) Model

SCR is developed with spatial factor model in connection with Gaussian processes, and variational inference is used to estimate spatially varying marginal dependence networks in a computationally efficient manner. We assume that $\boldsymbol{y}(\boldsymbol{s}_i) = (y_{i1}, \ldots, y_{ip})^\top \in \mathbb{R}^p$ denote the normalized and centered gene expression profiles for $p$ genes at spatial location $\boldsymbol{s}_i \in \boldsymbol{\mathcal{S}} \subset \mathbb{R}^2$. We aim to estimate spatially varying $p$-dimensional networks where the marginal dependencies between genes are captured by an undirected graph with a set of vertices $\mathbb{V} = \{1, \ldots, p\}$ and a set of spatially varying edges $\mathbb{E}(\boldsymbol{\mathcal{S}}) \subseteq \mathbb{V} \times \mathbb{V}$. In particular, we quantify the spatially varying co-expression level between two genes within the gene panel using the correlation coefficient.

#### S1.1 Model specification

##### S1.1.1 SCR model

The spatial covariance regression model is provided as follows wrt each spatial location $\boldsymbol{s}_i$

$$\boldsymbol{y}(\boldsymbol{s}_i) = \boldsymbol{\Lambda}(\boldsymbol{s}_i)\boldsymbol{\eta}_i + \boldsymbol{\varepsilon}_i, \quad \boldsymbol{\eta}_i \overset{\text{iid}}{\sim} \mathcal{N}_K(0, \mathbf{I}_K), \quad \boldsymbol{\varepsilon}_i \overset{\text{iid}}{\sim} \mathcal{N}_p(0, \boldsymbol{\Sigma}_0) \tag{S1}$$

where $\boldsymbol{\Lambda}(\boldsymbol{s}_i) \in \mathbb{R}^{p\times K}$ denotes the spatial factor loading matrix, $\boldsymbol{\eta}_i \in \mathbb{R}^K$ are the latent factor scores, and $\boldsymbol{\varepsilon}_i \in \mathbb{R}^p$ are zero-mean Gaussian errors with covariance $\boldsymbol{\Sigma}_0 \in \mathbb{R}^{p\times p}$. We assume $\boldsymbol{\Sigma}_0 = \text{diag}(\sigma_1^2, \ldots, \sigma_p^2)$ is diagonal and captures gene-specific residual variability. A low-dimensional structure is induced by assuming $K \ll p$.

**Charecterization of sptially covariance matrices.** By allowing the loadings to vary spatially, we enable the estimation of spatially varying covariance and correlation matrices.

$$\begin{aligned}
\text{Cov}(\boldsymbol{y}(\boldsymbol{s}_i)) &= \mathbb{E}\left[(\boldsymbol{\Lambda}(\boldsymbol{s}_i)\boldsymbol{\eta}_i + \boldsymbol{\varepsilon}_i)(\boldsymbol{\Lambda}(\boldsymbol{s}_i)\boldsymbol{\eta}_i + \boldsymbol{\varepsilon}_i)^\top\right] \\
&= \boldsymbol{\Lambda}(\boldsymbol{s}_i)\mathbb{E}\left[\boldsymbol{\eta}_i\boldsymbol{\eta}_i^\top\right]\boldsymbol{\Lambda}(\boldsymbol{s}_i)^\top + \mathbb{E}\left[\boldsymbol{\varepsilon}_i\boldsymbol{\varepsilon}_i^\top\right] + \mathbb{E}\left[\boldsymbol{\Lambda}(\boldsymbol{s}_i)\boldsymbol{\eta}_i\boldsymbol{\varepsilon}_i^\top\right] + \mathbb{E}\left[\boldsymbol{\varepsilon}_i\boldsymbol{\eta}_i^\top\boldsymbol{\Lambda}(\boldsymbol{s}_i)^\top\right] \\
&= \boldsymbol{\Lambda}(\boldsymbol{s}_i)\mathbf{I}_K\boldsymbol{\Lambda}(\boldsymbol{s}_i)^\top + \boldsymbol{\Sigma}_0 \\
&= \boldsymbol{\Lambda}(\boldsymbol{s}_i)\boldsymbol{\Lambda}(\boldsymbol{s}_i)^\top + \boldsymbol{\Sigma}_0
\end{aligned} \tag{S2}$$

##### S1.1.2 Identifiability

Factor models are invariant under orthogonal rotations of the factor loadings. In particular, for any orthogonal matrix $\mathbf{Q} \in \mathbb{R}^{K\times K}$, the transformation

$$\boldsymbol{\Lambda}(\boldsymbol{s}_i) \to \boldsymbol{\Lambda}(\boldsymbol{s}_i)\mathbf{Q}$$

leaves the marginal likelihood unchanged. Although the factor loadings are identifiable only up to a rotation, the covariance structure they encode is invariant to this non-identifiability. Since our primary interest lies in estimating the covariance and correlation structure, this rotational non-identifiability does not affect inference on the spatial gene networks. Concretely, let $\boldsymbol{\Lambda}^*(\boldsymbol{s}_i) \coloneqq \boldsymbol{\Lambda}(\boldsymbol{s}_i)\mathbf{Q}$. Since $\mathbf{Q}$ is orthogonal, $\mathbf{Q}\mathbf{Q}^\top = \mathbf{Q}^\top\mathbf{Q} = \mathbf{I}_K$. From Equation S2,

$$\begin{aligned}
\mathrm{Cov}^*(\boldsymbol{y}(\boldsymbol{s}_i)) &= \boldsymbol{\Lambda}^*(\boldsymbol{s}_i)\boldsymbol{\Lambda}^*(\boldsymbol{s}_i)^\top + \boldsymbol{\Sigma}_0 \\
&= \boldsymbol{\Lambda}(\boldsymbol{s}_i)\mathbf{Q}\mathbf{Q}^\top\boldsymbol{\Lambda}(\boldsymbol{s}_i)^\top + \boldsymbol{\Sigma}_0 \\
&= \boldsymbol{\Lambda}(\boldsymbol{s}_i)\boldsymbol{\Lambda}(\boldsymbol{s}_i)^\top + \boldsymbol{\Sigma}_0 \\
&= \mathrm{Cov}(\boldsymbol{y}(\boldsymbol{s}_i)).
\end{aligned}$$

Accordingly, even though the loading matrix is identifiable only up to an orthogonal rotation, $\mathrm{Cov}(\boldsymbol{y}(\boldsymbol{s}_i))$ is unaffected by this rotational non-identifiability.

#### S1.2 Variational inference

Bayesian inference for factor models is often carried out with Markov chain Monte Carlo (MCMC), but such algorithms scale poorly in high dimensions Lopes and West (2004). For scalability, we develop a coordinate ascent variational inference (CAVI) algorithm for estimation and uncertainty quantification of the model parameters

$$\boldsymbol{z} = \Big(\{\boldsymbol{\eta}_i\}_{i=1}^n,\ \{\boldsymbol{\theta}_g\}_{g=1}^p,\ \{\phi_{gr}\}_{g,r},\ \{\delta_r\}_{r=1}^L,\ \{\boldsymbol{\xi}_{rk}\}_{r,k},\ \{\sigma_g^2\}_{g=1}^p\Big),$$

where $\boldsymbol{\theta}_g = (\theta_{g1}, \ldots, \theta_{gL})^\top \in \mathbb{R}^L$ denotes row $g$ of $\boldsymbol{\Theta}$. Variational inference posits a family $\mathcal{Q}$ of densities over $\boldsymbol{z}$ and seeks the surrogate posterior $q(\boldsymbol{z}) \in \mathcal{Q}$ minimizing the Kullback–Leibler divergence to the true posterior,

$$\operatorname*{argmin}_{q(\boldsymbol{z})\in\mathcal{Q}} \mathrm{KL}\left(q(\boldsymbol{z}) \| p(\boldsymbol{z} \mid \boldsymbol{Y})\right),$$

equivalently maximizing the evidence lower bound (ELBO),

$$\operatorname*{argmax}_{q(\boldsymbol{z})\in\mathcal{Q}} \mathrm{ELBO}(q) = \operatorname*{argmax}_{q(\boldsymbol{z})\in\mathcal{Q}} \left\{\mathbb{E}_q\left[\log p(\boldsymbol{z}, \boldsymbol{Y})\right] - \mathbb{E}_q\left[\log q(\boldsymbol{z})\right]\right\}.$$

We adopt a mean-field family that partitions the parameters into disjoint blocks Hansen et al. (2025). Letting $\boldsymbol{z}_{-b}$ denote all blocks except $\boldsymbol{z}_b$, the optimal factor satisfies

$$q_b(\boldsymbol{z}_b) \propto \exp\Big(\mathbb{E}_{q(\boldsymbol{z}_{-b})}\left[\log p(\boldsymbol{z}_b \mid \boldsymbol{z}_{-b}, \boldsymbol{Y})\right]\Big).$$

When the complete conditional belongs to the exponential family, the optimal factor remains in the same family, reducing ELBO maximization to updating the associated natural parameters. Under the mean-field assumption the variational posterior factorizes as

$$\begin{aligned}
q(\boldsymbol{z}) = &\left[\prod_{i=1}^n q\left(\boldsymbol{\eta}_i \mid \boldsymbol{\mu}_i^\eta, \boldsymbol{\Sigma}_i^\eta\right)\right]\left[\prod_{g=1}^p q\Big(\boldsymbol{\theta}_g \mid \boldsymbol{\mu}_g^\Theta, \boldsymbol{\Sigma}_g^\Theta\Big)\right]\left[\prod_{g=1}^p\prod_{r=1}^L q\Big(\phi_{gr} \mid a_{gr}^\phi, b_{gr}^\phi\Big)\right] \\
&\times \left[\prod_{r=1}^L q\Big(\delta_r \mid a_r^\delta, b_r^\delta\Big)\right]\left[\prod_{r=1}^L\prod_{k=1}^K q\left(\boldsymbol{\xi}_{rk} \mid \boldsymbol{m}_{rk}, \mathbf{S}_{rk}\right)\right]\left[\prod_{g=1}^p q\Big(\sigma_g^{-2} \mid a_g^\sigma, b_g^\sigma\Big)\right].
\end{aligned}$$

We write $\mu_{gr}^\theta = [\boldsymbol{\mu}_g^\Theta]_r$ and $v_{gr}^\theta = [\boldsymbol{\Sigma}_g^\Theta]_{rr}$ for the marginal mean and variance of $\theta_{gr}$; $\mu_{ik}^\eta = [\boldsymbol{\mu}_i^\eta]_k$ and $v_{ik}^\eta = [\boldsymbol{\Sigma}_i^\eta]_{kk}$ for those of $\eta_{ik}$; and $\mu_{i,rk}^\Xi = [\boldsymbol{m}_{rk}]_i$ and $v_{i,rk}^\Xi = [\mathbf{S}_{rk}]_{ii}$ for those of $\Xi_{i,rk}$. Some algebraic steps below are omitted for brevity.

##### S1.2.1 Update for $\boldsymbol{\eta}_i$

Writing $\boldsymbol{\Xi}_i \coloneqq \boldsymbol{\Xi}(\boldsymbol{s}_i)$, the relevant terms are

$$\boldsymbol{y}_i = \boldsymbol{\Theta}\boldsymbol{\Xi}_i\boldsymbol{\eta}_i + \boldsymbol{\varepsilon}_i, \qquad \boldsymbol{\varepsilon}_i \sim \mathcal{N}_p(\mathbf{0}, \boldsymbol{\Sigma}_0), \qquad \boldsymbol{\eta}_i \sim \mathcal{N}_K(\mathbf{0}, \mathbf{I}_K).$$

The conditional log-likelihood, up to constants, is $-\frac{1}{2}(\boldsymbol{y}_i - \boldsymbol{\Theta}\boldsymbol{\Xi}_i\boldsymbol{\eta}_i)^\top \boldsymbol{\Sigma}_0^{-1}(\boldsymbol{y}_i - \boldsymbol{\Theta}\boldsymbol{\Xi}_i\boldsymbol{\eta}_i)$, which expands to

$$\boldsymbol{y}_i^\top \boldsymbol{\Sigma}_0^{-1}\boldsymbol{y}_i - 2\boldsymbol{y}_i^\top \boldsymbol{\Sigma}_0^{-1}\boldsymbol{\Theta}\boldsymbol{\Xi}_i\boldsymbol{\eta}_i + \boldsymbol{\eta}_i^\top \boldsymbol{\Xi}_i^\top \boldsymbol{\Theta}^\top \boldsymbol{\Sigma}_0^{-1}\boldsymbol{\Theta}\boldsymbol{\Xi}_i\boldsymbol{\eta}_i.$$

Adding the prior contribution $-\frac{1}{2}\boldsymbol{\eta}_i^\top \boldsymbol{\eta}_i$ and taking variational expectations over $\boldsymbol{\Theta}, \boldsymbol{\Xi}_i, \boldsymbol{\Sigma}_0$ gives

$$\log q^*(\boldsymbol{\eta}_i) = -\tfrac{1}{2}\boldsymbol{\eta}_i^\top \Big(\mathbf{I}_K + \mathbb{E}[\boldsymbol{\Xi}_i^\top \boldsymbol{\Psi}\boldsymbol{\Xi}_i]\Big)\boldsymbol{\eta}_i + \boldsymbol{\eta}_i^\top \mathbb{E}[\boldsymbol{\Xi}_i^\top]\mathbb{E}[\boldsymbol{\Theta}^\top]\mathbb{E}[\boldsymbol{\Sigma}_0^{-1}]\boldsymbol{y}_i + \text{const},$$

where

$$\boldsymbol{\Psi} \coloneqq \mathbb{E}[\boldsymbol{\Theta}^\top \boldsymbol{\Sigma}_0^{-1}\boldsymbol{\Theta}] = \sum_{g=1}^{p} \frac{a_g^\sigma}{b_g^\sigma}\Big(\boldsymbol{\mu}_g^\Theta(\boldsymbol{\mu}_g^\Theta)^\top + \boldsymbol{\Sigma}_g^\Theta\Big) \in \mathbb{R}^{L\times L}.$$

Hence $q^*(\boldsymbol{\eta}_i) = \mathcal{N}_K(\boldsymbol{\mu}_i^\eta, \boldsymbol{\Sigma}_i^\eta)$ with

$$\boldsymbol{\Sigma}_i^\eta = \Big(\mathbf{I}_K + \mathbb{E}[\boldsymbol{\Xi}_i^\top \boldsymbol{\Psi}\boldsymbol{\Xi}_i]\Big)^{-1}, \qquad \boldsymbol{\mu}_i^\eta = \boldsymbol{\Sigma}_i^\eta \mathbb{E}[\boldsymbol{\Xi}_i^\top]\mathbb{E}[\boldsymbol{\Theta}^\top]\mathbb{E}[\boldsymbol{\Sigma}_0^{-1}]\boldsymbol{y}_i.$$

Since the columns of $\boldsymbol{\Xi}_i$ have diagonal variational covariance $\boldsymbol{\Sigma}_{ik}^\Xi = \operatorname{diag}(v_{i,1k}^\Xi, \dots, v_{i,Lk}^\Xi)$, the required expectation is, elementwise,

$$\big(\mathbb{E}[\boldsymbol{\Xi}_i^\top \boldsymbol{\Psi}\boldsymbol{\Xi}_i]\big)_{kk'} = (\boldsymbol{\mu}_{ik}^\Xi)^\top \boldsymbol{\Psi}\boldsymbol{\mu}_{ik'}^\Xi + \delta_{kk'}\sum_{r=1}^{L} \Psi_{rr} v_{i,rk}^\Xi,$$

where $\boldsymbol{\mu}_{ik}^\Xi = (\mu_{i,1k}^\Xi, \dots, \mu_{i,Lk}^\Xi)^\top \in \mathbb{R}^L$ is the $k$th column of $\mathbb{E}[\boldsymbol{\Xi}_i]$ and $\delta_{kk'}$ is the Kronecker delta.

##### S1.2.2 Update for $\boldsymbol{\theta}_g$ (row $g$ of $\boldsymbol{\Theta}$)

From the elementwise model $y_{ig} = \boldsymbol{\theta}_g^\top \boldsymbol{\Xi}_i\boldsymbol{\eta}_i + \varepsilon_{ig}$, $\varepsilon_{ig} \sim \mathcal{N}(0, \sigma_g^2)$, with multiplicative gamma process shrinkage prior $\theta_{gr} \mid \phi_{gr}, \tau_r \sim \mathcal{N}(0, (\phi_{gr}\tau_r)^{-1})$, the log-prior is $-\frac{1}{2}\sum_{r=1}^{L}\phi_{gr}\tau_r\theta_{gr}^2 + \text{const}$ and the conditional log-likelihood is $-\frac{1}{2}\sigma_g^{-2}\sum_{i=1}^{n}(y_{ig} - \boldsymbol{\theta}_g^\top \boldsymbol{\Xi}_i\boldsymbol{\eta}_i)^2$. Expanding each square and dropping terms free of $\boldsymbol{\theta}_g$,

$$\log q^*(\boldsymbol{\theta}_g) = -\tfrac{1}{2}\boldsymbol{\theta}_g^\top \Big(\mathbf{D}_g + \sigma_g^{-2}\sum_{i=1}^{n}\boldsymbol{\Xi}_i\boldsymbol{\eta}_i\boldsymbol{\eta}_i^\top \boldsymbol{\Xi}_i^\top\Big)\boldsymbol{\theta}_g + \sigma_g^{-2}\boldsymbol{\theta}_g^\top \sum_{i=1}^{n} y_{ig}\boldsymbol{\Xi}_i\boldsymbol{\eta}_i + \text{const},$$

with $\mathbf{D}_g = \operatorname{diag}(\phi_{g1}\tau_1, \dots, \phi_{gL}\tau_L)$. Collecting the $n$ locations through

$$\boldsymbol{\Gamma} \coloneqq \begin{bmatrix} (\boldsymbol{\Xi}_1\boldsymbol{\eta}_1)^\top \\ \vdots \\ (\boldsymbol{\Xi}_n\boldsymbol{\eta}_n)^\top \end{bmatrix} \in \mathbb{R}^{n\times L}, \qquad \boldsymbol{y}_g = (y_{1g}, \dots, y_{ng})^\top \in \mathbb{R}^n,$$

so that $\sum_i \boldsymbol{\Xi}_i\boldsymbol{\eta}_i\boldsymbol{\eta}_i^\top \boldsymbol{\Xi}_i^\top = \boldsymbol{\Gamma}^\top \boldsymbol{\Gamma}$ and $\sum_i y_{ig}\boldsymbol{\Xi}_i\boldsymbol{\eta}_i = \boldsymbol{\Gamma}^\top \boldsymbol{y}_g$, and taking expectations yields the joint Gaussian $q^*(\boldsymbol{\theta}_g) = \mathcal{N}_L(\boldsymbol{\mu}_g^\Theta, \boldsymbol{\Sigma}_g^\Theta)$ with full covariance

$$\boldsymbol{\Sigma}_g^{\Theta} = \Big(\mathbb{E}[\mathbf{D}_g] + \mathbb{E}[\sigma_g^{-2}]\mathbb{E}[\boldsymbol{\Gamma}^\top\boldsymbol{\Gamma}]\Big)^{-1}, \qquad \boldsymbol{\mu}_g^{\Theta} = \mathbb{E}[\sigma_g^{-2}]\boldsymbol{\Sigma}_g^{\Theta}\mathbb{E}[\boldsymbol{\Gamma}]^\top \boldsymbol{y}_g,$$

where

$$\mathbb{E}[\sigma_g^{-2}] = \frac{a_g^\sigma}{b_g^\sigma}, \qquad \mathbb{E}[\mathbf{D}_g] = \operatorname{diag}\left(\frac{a_{g1}^\phi}{b_{g1}^\phi}\cdot\frac{a_1^\delta}{b_1^\delta},\ \ldots,\ \frac{a_{gL}^\phi}{b_{gL}^\phi}\cdot\prod_{\ell=1}^{L}\frac{a_\ell^\delta}{b_\ell^\delta}\right).$$

The moments of $\boldsymbol{\Gamma}$ are

$$\mathbb{E}[\boldsymbol{\Gamma}] = \begin{bmatrix}\mathbb{E}[\boldsymbol{\Xi}_1]\boldsymbol{\mu}_1^\eta \\ \vdots \\ \mathbb{E}[\boldsymbol{\Xi}_n]\boldsymbol{\mu}_n^\eta\end{bmatrix}, \qquad \mathbb{E}[\boldsymbol{\Gamma}^\top\boldsymbol{\Gamma}] = \sum_{i=1}^{n}\mathbb{E}[\boldsymbol{\Xi}_i\boldsymbol{\Omega}_i^\eta\boldsymbol{\Xi}_i^\top], \quad \boldsymbol{\Omega}_i^\eta = \boldsymbol{\Sigma}_i^\eta + \boldsymbol{\mu}_i^\eta(\boldsymbol{\mu}_i^\eta)^\top,$$

where $\boldsymbol{\Omega}_i^\eta$ is the variational second moment of $\boldsymbol{\eta}_i$. With the diagonal column covariances $\boldsymbol{\Sigma}_{ik}^{\Xi} = \operatorname{diag}(v_{i,1k}^{\Xi}, \ldots, v_{i,Lk}^{\Xi})$,

$$\mathbb{E}[\boldsymbol{\Xi}_i\boldsymbol{\Omega}_i^\eta\boldsymbol{\Xi}_i^\top] = \sum_{k=1}^{K}\sum_{k'=1}^{K}(\boldsymbol{\Omega}_i^\eta)_{kk'}\Big(\boldsymbol{\mu}_{ik}^{\Xi}(\boldsymbol{\mu}_{ik'}^{\Xi})^\top + \delta_{kk'}\boldsymbol{\Sigma}_{ik}^{\Xi}\Big).$$

The rows of $\boldsymbol{\Theta}$ form the variational blocks: each $\boldsymbol{\theta}_g$ is updated jointly with the full $L \times L$ covariance $\boldsymbol{\Sigma}_g^{\Theta}$, retaining the within-row posterior correlations rather than only its diagonal.

##### S1.2.3 Update for local shrinkage $\phi_{gr}$

With $\theta_{gr} \mid \phi_{gr}, \tau_r \sim \mathcal{N}(0, (\phi_{gr}\tau_r)^{-1})$ and $\phi_{gr} \sim \mathrm{Ga}(\nu/2, \nu/2)$,

$$\begin{aligned}\log p(\phi_{gr} \mid -) &= \Big(\tfrac{\nu}{2} - 1\Big)\log\phi_{gr} - \tfrac{\nu}{2}\phi_{gr} + \tfrac{1}{2}\log(\phi_{gr}\tau_r) - \tfrac{1}{2}\phi_{gr}\tau_r\theta_{gr}^2 + \text{const} \\ &= \Big(\tfrac{\nu+1}{2} - 1\Big)\log\phi_{gr} - \tfrac{1}{2}(\nu + \tau_r\theta_{gr}^2)\phi_{gr} + \text{const}.\end{aligned}$$

Taking expectations, $q^*(\phi_{gr}) = \mathrm{Ga}(a_{gr}^\phi, b_{gr}^\phi)$ with

$$a_{gr}^\phi = \frac{\nu+1}{2}, \qquad b_{gr}^\phi = \tfrac{1}{2}\big[\nu + \mathbb{E}[\tau_r]\mathbb{E}[\theta_{gr}^2]\big],$$

where $\mathbb{E}[\tau_r] = \prod_{\ell=1}^{r}\frac{a_\ell^\delta}{b_\ell^\delta}$ and $\mathbb{E}[\theta_{gr}^2] = (\mu_{gr}^\theta)^2 + [\boldsymbol{\Sigma}_g^{\Theta}]_{rr}$.

##### S1.2.4 Update for global shrinkage $\delta_r$

Recall $\tau_r = \prod_{\ell=1}^{r}\delta_\ell$, with $\delta_1 \sim \mathrm{Ga}(a_1, 1)$ and $\delta_r \sim \mathrm{Ga}(a_2, 1)$ for $r \geq 2$; only $\{\theta_{gr'} : r' \geq r\}$ depend on $\delta_r$. Collecting the $\delta_r$-dependent terms,

$$\log p(\delta_r \mid -) = \Big(a_r - 1 + \tfrac{p(L-r+1)}{2}\Big)\log\delta_r - \delta_r\Big[1 + \tfrac{1}{2}\sum_{g=1}^{p}\sum_{r'=r}^{L}\phi_{gr'}\theta_{gr'}^2\prod_{\substack{\ell=1\\ \ell\neq r}}^{r'}\delta_\ell\Big] + \text{const},$$

where $a_r = a_1$ if $r = 1$ and $a_r = a_2$ otherwise. Taking expectations, $q^*(\delta_r) = \mathrm{Ga}(a_r^\delta, b_r^\delta)$ with $a_r^\delta = a_r + \frac{p(L-r+1)}{2}$ and

$$b^{\delta}_{r} = 1 + \tfrac{1}{2}\sum_{g=1}^{p}\sum_{r'=r}^{L}\frac{a^{\phi}_{gr'}}{b^{\phi}_{gr'}}\Big((\mu^{\theta}_{gr'})^{2} + [\boldsymbol{\Sigma}^{\Theta}_{g}]_{r'r'}\Big)\prod_{\substack{\ell=1\\ \ell\neq r}}^{r'}\frac{a^{\delta}_{\ell}}{b^{\delta}_{\ell}}.$$

##### S1.2.5 Update for residual precisions $\sigma_g^{-2}$

With prior $\sigma_g^{-2} \sim \mathrm{Ga}(a_0, b_0)$ and likelihood $y_{ig} \sim \mathcal{N}(\boldsymbol{\theta}_g^{\top}\boldsymbol{\Xi}_i\boldsymbol{\eta}_i, \sigma_g^2)$, we obtain $q^*(\sigma_g^{-2}) = \mathrm{Ga}(a_g^{\sigma}, b_g^{\sigma})$ with $a_g^{\sigma} = a_0 + \frac{n}{2}$ and

$$b^{\sigma}_{g} = b_0 + \tfrac{1}{2}\sum_{i=1}^{n}\Big\{y_{ig}^{2} - 2y_{ig}(\boldsymbol{\mu}^{\Theta}_{g})^{\top}\sum_{k=1}^{K}\boldsymbol{\mu}^{\Xi}_{ik}\mu^{\eta}_{ik} + \mathrm{tr}\left[(\boldsymbol{\mu}^{\Theta}_{g}(\boldsymbol{\mu}^{\Theta}_{g})^{\top} + \boldsymbol{\Sigma}^{\Theta}_{g})\mathbf{G}_i\right]\Big\},$$

where $\boldsymbol{\mu}_i^{\Xi} \in \mathbb{R}^{L\times K}$ is $\mathbb{E}[\boldsymbol{\Xi}_i]$ with columns $\boldsymbol{\mu}^{\Xi}_{ik}$, and

$$\boldsymbol{\Omega}^{\eta}_{i} = \boldsymbol{\Sigma}^{\eta}_{i} + \boldsymbol{\mu}^{\eta}_{i}(\boldsymbol{\mu}^{\eta}_{i})^{\top}, \qquad \mathbf{G}_i = \boldsymbol{\mu}^{\Xi}_{i}\boldsymbol{\Omega}^{\eta}_{i}(\boldsymbol{\mu}^{\Xi}_{i})^{\top} + \sum_{k=1}^{K}(\boldsymbol{\Omega}^{\eta}_{i})_{kk}\boldsymbol{\Sigma}^{\Xi}_{ik}.$$

##### S1.2.6 Update for GP vectors $\boldsymbol{\xi}_{rk}$

Fix $(r,k)$. For each $(i,g)$,

$$y_{ig} = \underbrace{\theta_{gr}\eta_{ik}\xi_{rk}(\boldsymbol{s}_i)}_{\text{depends on } \boldsymbol{\xi}_{rk}} + \sum_{(r',k')\neq(r,k)}\theta_{gr'}\eta_{ik'}\xi_{r'k'}(\boldsymbol{s}_i) + \varepsilon_{ig}, \qquad \varepsilon_{ig} \sim \mathcal{N}(0,\sigma_g^2),$$

with blockwise partial residual

$$\widetilde{y}_{ig}^{-(r,k)} = y_{ig} - \sum_{(r',k')\neq(r,k)}\theta_{gr'}\eta_{ik'}\xi_{r'k'}(\boldsymbol{s}_i).$$

Summing the Gaussian log-likelihood over $g = 1,\ldots,p$ and discarding terms free of $\boldsymbol{\xi}_{rk}$,

$$\log p(\boldsymbol{Y} \mid -) = -\tfrac{1}{2}\sum_{i=1}^{n}\underbrace{\Big(\sum_{g=1}^{p}\sigma_g^{-2}\theta_{gr}^{2}\eta_{ik}^{2}\Big)}_{\widetilde{\alpha}_{i,rk}}\big(\xi_{rk}(\boldsymbol{s}_i)\big)^{2} + \sum_{i=1}^{n}\underbrace{\Big(\sum_{g=1}^{p}\sigma_g^{-2}\theta_{gr}\eta_{ik}\widetilde{y}_{ig}^{-(r,k)}\Big)}_{\widetilde{\beta}_{i,rk}}\xi_{rk}(\boldsymbol{s}_i) + \text{const}.$$

Adding the GP prior $\boldsymbol{\xi}_{rk} \sim \mathcal{N}_n(\mathbf{0},\mathbf{H})$, which contributes $-\frac{1}{2}\boldsymbol{\xi}_{rk}^{\top}\mathbf{H}^{-1}\boldsymbol{\xi}_{rk}$,

$$\log p(\boldsymbol{Y},\boldsymbol{\xi}_{rk} \mid -) = -\tfrac{1}{2}\boldsymbol{\xi}_{rk}^{\top}(\mathbf{H}^{-1} + \widetilde{\mathbf{D}}_{rk})\boldsymbol{\xi}_{rk} + \boldsymbol{\xi}_{rk}^{\top}\widetilde{\mathbf{b}}_{rk} + \text{const},$$

with $\widetilde{\mathbf{D}}_{rk} = \mathrm{diag}(\widetilde{\alpha}_{1,rk},\ldots,\widetilde{\alpha}_{n,rk})$ and $\widetilde{\mathbf{b}}_{rk} = (\widetilde{\beta}_{1,rk},\ldots,\widetilde{\beta}_{n,rk})^{\top}$. The mean-field update maximizes $\mathbb{E}[\log p(\boldsymbol{Y},\boldsymbol{\xi}_{rk} \mid -)]$ over $q(\boldsymbol{\xi}_{rk})$ with the other factors held fixed. Taking expectations,

$$\mathbb{E}[\widetilde{\alpha}_{i,rk}] = \sum_{g=1}^{p}\mathbb{E}[\sigma_g^{-2}]\mathbb{E}[\theta_{gr}^{2}]\mathbb{E}[\eta_{ik}^{2}] =: \alpha_{i,rk},$$

$$\mathbb{E}[\widetilde{\beta}_{i,rk}] = \sum_{g=1}^{p}\mathbb{E}[\sigma_g^{-2}]\mathbb{E}[\theta_{gr}]\mathbb{E}[\eta_{ik}]\bar{y}_{ig}^{-(r,k)} =: \beta_{i,rk},$$

where, under the factorization, the expected partial residual is

$$\bar{y}_{ig}^{-(r,k)} = y_{ig} - \sum_{(r',k')\neq(r,k)} \mu^{\theta}_{gr'}\mu^{\eta}_{ik'}\mu^{\Xi}_{i,r'k'}.$$

Substituting the variational moments $\mathbb{E}[\sigma_g^{-2}] = a_g^{\sigma}/b_g^{\sigma}$, $\mathbb{E}[\theta_{gr}] = \mu^{\theta}_{gr}$, $\mathbb{E}[\theta_{gr}^2] = (\mu^{\theta}_{gr})^2 + v^{\theta}_{gr}$, $\mathbb{E}[\eta_{ik}] = \mu^{\eta}_{ik}$, $\mathbb{E}[\eta_{ik}^2] = (\mu^{\eta}_{ik})^2 + v^{\eta}_{ik}$, and $\mathbb{E}[\xi_{r'k'}(\boldsymbol{s}_i)] = \mu^{\Xi}_{i,r'k'}$,

$$\alpha_{i,rk} = \sum_{g=1}^{p} \frac{a_g^{\sigma}}{b_g^{\sigma}}\big((\mu^{\theta}_{gr})^2 + v^{\theta}_{gr}\big)\big((\mu^{\eta}_{ik})^2 + v^{\eta}_{ik}\big), \qquad \beta_{i,rk} = \sum_{g=1}^{p} \frac{a_g^{\sigma}}{b_g^{\sigma}}\mu^{\theta}_{gr}\mu^{\eta}_{ik}\bar{y}_{ig}^{-(r,k)}.$$

With $\mathbf{D}_{rk} = \mathrm{diag}(\alpha_{1,rk}, \ldots, \alpha_{n,rk})$ and $\mathbf{b}_{rk} = (\beta_{1,rk}, \ldots, \beta_{n,rk})^{\top}$, the update is

$$q^*(\boldsymbol{\xi}_{rk}) = \mathcal{N}_n(\boldsymbol{m}_{rk}, \mathbf{S}_{rk}), \qquad \mathbf{S}_{rk}^{-1} = \mathbf{H}^{-1} + \mathbf{D}_{rk}, \qquad \boldsymbol{m}_{rk} = \mathbf{S}_{rk}\mathbf{b}_{rk},$$

from which we expose $\mu^{\Xi}_{i,rk} = [\boldsymbol{m}_{rk}]_i$ and $v^{\Xi}_{i,rk} = [\mathbf{S}_{rk}]_{ii}$.

##### S1.2.7 Initialization

Because CAVI converges only to a local optimum of the ELBO, a sensible initialization is important. We use a three-stage heuristic that seeds the variational means from a sparse principal component decomposition, reconstructs the spatial bases by per-spot least squares, and then performs a single mutually consistent refinement Hansen et al. (2025).

**Stage 1: Sparse PCA.** We apply sparse PCA with $L$ components to the data $\boldsymbol{Y} \in \mathbb{R}^{n\times p}$, obtaining a loading matrix $\boldsymbol{\Theta} \in \mathbb{R}^{p\times L}$ and scores $\mathbf{F} \in \mathbb{R}^{n\times L}$. Each score column is standardized to unit variance and the scaling absorbed into the corresponding loading column,

$$s_r = \mathrm{sd}(\mathbf{F}_{\cdot r}), \qquad \mathbf{F}_{\cdot r} \leftarrow \mathbf{F}_{\cdot r}/s_r, \qquad \boldsymbol{\Theta}_{\cdot r} \leftarrow s_r\boldsymbol{\Theta}_{\cdot r},$$

which leaves the reconstruction $\mathbf{F}\boldsymbol{\Theta}^{\top}$ unchanged. The first $K$ score columns initialize the factor means, $\boldsymbol{\mu}_i^{\eta} = (F_{i1}, \ldots, F_{iK})^{\top}$ (requiring $K \leq L$), and $\boldsymbol{\Theta}$ initializes the loading means $\{\boldsymbol{\mu}_g^{\Theta}\}$.

**Stage 2: Per-spot least squares for $\boldsymbol{\Xi}_i$ and noise.** Holding $\boldsymbol{\Theta}$ and the factor scores fixed, we reconstruct each local basis matrix from the relation $\boldsymbol{\Lambda}(\boldsymbol{s}_i)\boldsymbol{\eta}_i = \boldsymbol{\Theta}\boldsymbol{\Xi}_i\boldsymbol{\eta}_i \approx \boldsymbol{y}_i$. Regressing $\boldsymbol{y}_i$ on $\boldsymbol{\Theta}$ gives

$$\boldsymbol{w}_i = (\boldsymbol{\Theta}^{\top}\boldsymbol{\Theta})^{-1}\boldsymbol{\Theta}^{\top}\boldsymbol{y}_i \in \mathbb{R}^L,$$

and the minimum-norm $\boldsymbol{\Xi}_i$ satisfying $\boldsymbol{\Xi}_i\boldsymbol{\eta}_i = \boldsymbol{w}_i$ is the rank-one matrix

$$\boldsymbol{\Xi}_i = \frac{\boldsymbol{w}_i\boldsymbol{\eta}_i^{\top}}{\|\boldsymbol{\eta}_i\|^2}, \qquad \mu^{\Xi}_{i,rk} = [\boldsymbol{\Xi}_i]_{rk}.$$

The regression residual $\boldsymbol{\rho}_i = \boldsymbol{y}_i - \boldsymbol{\Theta}\boldsymbol{w}_i$ yields a per-spot variance $\hat{\sigma}_i^2 = \|\boldsymbol{\rho}_i\|^2/\max(p - \mathrm{rank}(\boldsymbol{\Theta}), 1)$, and propagating its uncertainty through the linear map seeds the basis variances,

$$\mathrm{Var}(\boldsymbol{w}_i) = \hat{\sigma}_i^2\mathrm{diag}((\boldsymbol{\Theta}^{\top}\boldsymbol{\Theta})^{-1}), \qquad v^{\Xi}_{i,rk} = \mathrm{Var}(w_{ir})\frac{\eta_{ik}^2}{\|\boldsymbol{\eta}_i\|^4}.$$

The residual variances are initialized from the diagonal mismatch between the empirical covariance $\mathbf{C} = \widehat{\mathrm{Cov}}(\boldsymbol{Y}) \in \mathbb{R}^{p\times p}$ and the low-rank reconstruction, taken at the median over spots,

$$\hat{\sigma}_g^2 = \max\left( \underset{i=1,\dots,n}{\text{median}} \left[\mathbf{C} - \boldsymbol{\Lambda}(\boldsymbol{s}_i)\boldsymbol{\Lambda}(\boldsymbol{s}_i)^\top\right]_{gg},\ \epsilon \right), \qquad \boldsymbol{\Lambda}(\boldsymbol{s}_i) = \boldsymbol{\Theta}\boldsymbol{\Xi}_i,$$

with a small floor $\epsilon > 0$. These set the residual-precision factor through $a_g^\sigma = a_0 + n/2$ and $b_g^\sigma = a_g^\sigma \hat{\sigma}_g^2$, so that $\mathbb{E}[\sigma_g^{-2}] = 1/\hat{\sigma}_g^2$.

**Shrinkage and prior factors.** The local- and global-shrinkage factors are initialized at their priors: $a_{gr}^\phi = b_{gr}^\phi = \nu/2$, and $(a_r^\delta, b_r^\delta)$ are taken from the multiplicative-gamma hyperparameters (with shape $a_1$ for the first column and $a_2$ thereafter). The factor-score covariances are set to $\boldsymbol{\Sigma}_i^\eta = \mathbf{I}_K$, matching the prior precision.

**Stage 3: Consistency refinement.** Finally, we perform one update of the global loadings $\{\boldsymbol{\theta}_g\}$ followed by one update of the factor scores $\{\boldsymbol{\eta}_i\}$, using the block updates derived above, so that the seeded means and covariances are mutually consistent before the main CAVI iterations begin.

#### S1.3 Computational considerations

The update for each spatial basis vector $\boldsymbol{\xi}_{rk}$ yields a Gaussian posterior with covariance $\mathbf{S}_{rk} = (\mathbf{H}^{-1} + \mathbf{D}_{rk})^{-1}$, where $\mathbf{H} \in \mathbb{R}^{n \times n}$ is the kernel matrix with $(\mathbf{H})_{ij} = \kappa(\boldsymbol{s}_i, \boldsymbol{s}_j)$ and $\mathbf{D}_{rk}$ is diagonal and depends on the current variational expectations. Suppressing the $(r, k)$ subscript and writing $\mathbf{S} = (\mathbf{H}^{-1} + \mathbf{D})^{-1}$, the Woodbury identity gives

$$\mathbf{S} = \mathbf{H} - \mathbf{H}(\mathbf{H} + \mathbf{D}^{-1})^{-1}\mathbf{H}.$$

Let $\mathbf{W} := \mathbf{H} + \mathbf{D}^{-1}$. Computing $\text{diag}(\mathbf{S})$ exactly would require solving the linear system $\mathbf{W}\mathbf{X} = \mathbf{H}$ with $n$ right-hand sides, which is computationally expensive for large $n$. Since we only require the marginal posterior variances $\mathbf{S}_{ii}$, we estimate $\text{diag}(\mathbf{S})$ using Hutchinson's stochastic diagonal estimator Bekas et al. (2007). Let $\boldsymbol{z} \in \{-1, +1\}^n$ be a random Rademacher vector with independent entries. For any symmetric matrix $\mathbf{A}$,

$$\text{diag}(\mathbf{A}) = \mathbb{E}\left[\boldsymbol{z} \odot (\mathbf{A}\boldsymbol{z})\right], \tag{S3}$$

where $\odot$ and $\oslash$ denote elementwise product and division, respectively. Applying (S3) to $\mathbf{A} = \mathbf{S}$, and noting that

$$\mathbf{S}\boldsymbol{z} = \mathbf{H}\boldsymbol{z} - \mathbf{H}\mathbf{W}^{-1}(\mathbf{H}\boldsymbol{z}),$$

we approximate the diagonal by averaging over $J$ independent probe vectors $\{\boldsymbol{z}_t\}_{t=1}^J$:

$$\widehat{\text{diag}(\mathbf{S})} = \left[\sum_{t=1}^{J} \boldsymbol{z}_t \odot \mathbf{S}\boldsymbol{z}_t\right] \oslash \left[\sum_{t=1}^{J} \boldsymbol{z}_t \odot \boldsymbol{z}_t\right].$$

This avoids explicit construction of the dense matrix $\mathbf{S}$ and replaces an $n$-right-hand-side linear solve with $J \ll n$ right-hand sides. Concretely, instead of forming $\mathbf{S} = \mathbf{H} - \mathbf{H}\mathbf{W}^{-1}\mathbf{H}$ exactly, which requires solving against all $n$ columns of $\mathbf{H}$ and forming dense $n \times n$ products, we solve

$$\mathbf{W}\mathbf{Y} = \mathbf{H}\mathbf{Z}$$

with only the $J$ probe vectors collected in $\mathbf{Z}$, and estimate $\text{diag}(\mathbf{S})$ from the resulting quantities. Since these operations scale with $J$ rather than $n$, the cost and memory footprint are substantially smaller when $J \ll n$.

Table S1: Notation summary and indexing conventions.

| Symbol | Meaning | Info |
|---|---|---|
| $\boldsymbol{\mathcal{S}}$ | Spatial domain | $\subset \mathbb{R}^2$ |
| $\mathbf{s}_i$ | Spatial location $i$ | $i = 1, \dots, n$ |
| $n$ | Number of spatial locations | $\in \mathbb{N}$ |
| $p$ | Number of genes | $\in \mathbb{N}$ |
| $\mathbf{y}(\mathbf{s}_i)$ or $\mathbf{y}_i$ | Gene expression at location $i$ | $\in \mathbb{R}^p$ |
| $y_{ig}$ | Expression of gene $g$ at location $i$ | $i = 1, \dots, n;\ g = 1, \dots, p$ |
| $\mathbb{V}$ | Graph vertex set (genes) | $\{1, \dots, p\}$ |
| $\mathbb{E}(\boldsymbol{\mathcal{S}})$ | Spatially varying edge set | $\subseteq \mathbb{V} \times \mathbb{V}$ |
| $\mathbb{G}(\boldsymbol{\mathcal{S}})$ | Spatially varying undirected graph | $(\mathbb{V}, \mathbb{E}(\boldsymbol{\mathcal{S}}))$ |
| $\Sigma(\mathbf{s}_i)$ | Covariance at location $i$ | $\in \mathbb{R}^{p \times p}$ |
| $\Sigma_0$ | Residual covariance (diagonal) | $\mathrm{diag}(\sigma_1^2, \dots, \sigma_p^2)$ |
| $\sigma_g^2$ | Residual variance for gene $g$ | $g = 1, \dots, p$ |
| $K$ | Number of latent factors | $\in \mathbb{N}$ |
| $L$ | Number of spatial bases (global bases) | $\in \mathbb{N}$ |
| $\eta_i$ | Latent factor scores at location $i$ | $\in \mathbb{R}^K$ |
| $\varepsilon_i$ | Residual noise at location $i$ | $\in \mathbb{R}^p$ |
| $\Lambda(\mathbf{s}_i)$ | Spatial loading matrix | $\in \mathbb{R}^{p \times K}$ |
| $\Theta$ | Global loading/basis matrix | $\in \mathbb{R}^{p \times L}$ |
| $\theta_{gr}$ | Entry of $\Theta$ for gene $g$, basis $r$ | $g = 1, \dots, p;\ r = 1, \dots, L$ |
| $\Xi(\mathbf{s}_i)$ or $\Xi_i$ | Local spatial basis matrix at location $i$ | $\in \mathbb{R}^{L \times K}$ |
| $\xi_{rk}(\mathbf{s})$ | Spatial basis function (GP) for $(r, k)$ | $\mathbf{s} \in \boldsymbol{\mathcal{S}}$ |
| $\boldsymbol{\xi}_{rk}$ | Evaluations of $\xi_{rk}(\cdot)$ over all locations | $\in \mathbb{R}^n$ |
| $\Xi_{i,rk}$ | Entry of $\Xi_i$ for basis $r$ and factor $k$ | $i = 1, \dots, n;\ r = 1, \dots, L;\ k = 1, \dots, K$ |
| $M$ | Number of posterior MC draws for SCG identification | $\in \mathbb{N}$ |
| $\kappa(\mathbf{s}, \mathbf{s}')$ | GP kernel function | $\boldsymbol{\mathcal{S}} \times \boldsymbol{\mathcal{S}} \to \mathbb{R}$ |
| $\rho$ | Kernel length-scale parameter | $> 0$ |
| $\zeta$ | RQ kernel shape parameter | $> 0$ |
| $\mathbf{H}$ | Kernel matrix with $\mathbf{H}_{ij} = \kappa(\mathbf{s}_i, \mathbf{s}_j)$ | $\in \mathbb{R}^{n \times n}$ |
| $\mathbf{D}$ | Diagonal precision matrix in $\xi$ update | $\in \mathbb{R}^{n \times n}$ (diagonal) |
| $\mathbf{W}$ | Linear system matrix, $\mathbf{W} = \mathbf{H} + \mathbf{D}^{-1}$ | $\in \mathbb{R}^{n \times n}$ |
| $\mathbf{S}$ | Posterior covariance, $\mathbf{S} = (\mathbf{H}^{-1} + \mathbf{D})^{-1}$ | $\in \mathbb{R}^{n \times n}$ |
| $\mathrm{diag}(\mathbf{S})$ | Posterior marginal variances of $\boldsymbol{\xi}_{rk}$ | $\in \mathbb{R}^n$ |
| $J$ | Number of Hutchinson probe vectors | $\in \mathbb{N}$, typically $J \ll n$ |
| $z_t$ | Rademacher probe vector | $\in \{-1, +1\}^n,\ t = 1, \dots, J$ |
| $i$ (index) | Spatial location index | $1, \dots, n$ |
| $g$ (index) | Gene index | $1, \dots, p$ |
| $r$ (index) | Spatial basis index | $1, \dots, L$ |
| $k$ (index) | Latent factor index | $1, \dots, K$ |
| $\ell$ (index) | Shrinkage-prior product index, $\tau_r = \prod_{\ell=1}^{r} \delta_\ell$ | $1, \dots, r$ |
| $b$ (index) | Mean-field block index | $1, \dots, B$ |

### S2 Additional Simulation Results

We conducted a comprehensive simulation study to assess the statistical accuracy and computational efficiency of SCR, evaluating (i) recovery of spatially varying network structure, (ii) identification of SCGs, and (iii) scalability with respect to increasing spatial resolution. We considered spatial grids of 5,000, 10,000, 15,000, 20,000, 30,000, 40,000, and 50,000 locations with $p = 30$ genes, summarizing results over 20 independent replicates.

To assess recovery of spatially varying network structure, we generated data from a controlled spatial factor model with nonlinear spatial effects and sparse gene loadings. Spatial locations $\{\boldsymbol{s}_i\}_{i=1}^n$ were arranged on a grid over the domain $\boldsymbol{\mathcal{S}} = [-1, 1]^2$, where $\boldsymbol{s}_i = (x_i, y_i) \in \boldsymbol{\mathcal{S}}$ denotes the coordinate of the $i$th location. We generated $K_{\text{spatial}}$ spatially varying latent factor fields and $K_{\text{const}}$ spatially constant factor fields. To induce heterogeneous spatial structure, we randomly sampled $K_{\text{spatial}}$ functions from a library of 100 nonlinear basis functions, including linear, polynomial, radial, exponential, logistic, and hyperbolic forms. For each selected function $f_k(x, y)$, we defined the spatial factor field

$$B_k(\boldsymbol{s}_i) = \frac{f_k(x_i, y_i) - \overline{f_k}}{\text{sd}(f_k)},$$

so that each field $B_k$ had mean zero and unit variance across locations. We also generated $K_{\text{const}}$ constant latent factors $B_k(\boldsymbol{s}_i) = c_k$, $c_k \sim \mathcal{N}(0, 1)$, fixed across all locations. Let

$$\mathbf{B}(\boldsymbol{s}_i) \in \mathbb{R}^{K_{\text{spatial}} + K_{\text{const}}}$$

denote the combined vector of spatial and constant factor values at location $\boldsymbol{s}_i$. Gene loadings were constructed to induce sparse, heterogeneous covariance patterns: a fraction $f_{\text{spatial}}$ of genes were assigned nonzero loadings on one or more spatial factors, with coefficients drawn from $\mathcal{N}(0, 1)$; a fraction $f_{\text{const}}$ of genes were assigned loadings on constant factors, with coefficients drawn from $\mathcal{N}(0, 0.01)$, inducing weaker global correlation; and the remaining genes had zero loadings. Letting $\boldsymbol{\Theta}^{\text{sim}} \in \mathbb{R}^{p \times (K_{\text{spatial}} + K_{\text{const}})}$ denote the resulting loading matrix, we defined the location-specific loading matrix

$$\boldsymbol{\Lambda}(\boldsymbol{s}_i) = \boldsymbol{\Theta}^{\text{sim}} \text{diag}(\mathbf{B}(\boldsymbol{s}_i)),$$

the spatial covariance matrix $\boldsymbol{\Sigma}(\boldsymbol{s}_i) = \boldsymbol{\Lambda}(\boldsymbol{s}_i)\boldsymbol{\Lambda}(\boldsymbol{s}_i)^\top + \sigma^2 \mathbf{I}_p$, and generated observed expression as $\boldsymbol{y}(\boldsymbol{s}_i) \sim \mathcal{N}_p(0, \boldsymbol{\Sigma}(\boldsymbol{s}_i))$. Ground-truth SCG labels were determined by computing the spatial standard deviation of the true correlation field for each gene pair and applying Otsu's thresholding to these values; pairs exceeding the threshold were labeled SCGs and all others non-SCGs. In each simulation scenario and replicate, approximately 10% of edges were specified as SCGs.

Under these settings, SCR demonstrated consistently strong recovery of spatially varying network structure across all spatial resolutions (Table S3). The mean Matthews correlation coefficient (MCC) ranged from 0.920 at $n = 5{,}000$ to 0.942 at $n = 50{,}000$, remaining between 0.928 and 0.942 for $n \geq 10{,}000$. Variability across replicates was low (standard deviation between 0.029 and 0.062), and even the worst-case replicate exceeded 0.71 at the smallest grid and 0.82 for $n \geq 10{,}000$. Sensitivity and specificity were likewise high: mean true positive rates (TPR) ranged from 0.962 to 0.990, mean false positive rates (FPR) remained between 0.010 and 0.016, and true negative rates exceeded 0.983 in all settings. Because SCGs constituted a minority of all gene pairs (approximately 10% by construction), this is a substantially imbalanced classification problem, and the consistently high MCC values demonstrate reliable discrimination between true and spurious edges despite the imbalance.

We further evaluated recovery of spatially varying correlation structure using the RV coefficient Hansen et al. (2025), which measures concordance between estimated and true spatial correlation matrices at each location. Across all resolutions, the mean RV coefficient ranged from 0.984 to 0.990, with mean standard deviation across replicates below 0.012; maximum RV values were near

1 and even the minimum observed value remained above 0.74, indicating robust spatial covariance recovery throughout the domain. Posterior calibration was likewise stable across resolutions: the estimated posterior false discovery rate (FDR) had mean values between 0.089 and 0.092, remaining below the nominal 10% target, and the average edge selection rate varied modestly between 11% and 12%, consistent with the data-generating process in which roughly 10% of gene pairs were SCGs.

Computational performance scaled predictably with the number of spatial locations (Figure S13). Averaging the total runtime over the 20 replicates and over the 435 gene pairs, the mean per-edge runtime grew from 1.3 seconds at $n = 5{,}000$ to 40.2 seconds at $n = 50{,}000$ (2.8, 4.7, 7.1, 14.3, and 24.6 seconds at $n = 10{,}000$, 15,000, 20,000, 30,000, and 40,000, respectively), corresponding to a mean wall-clock time of roughly 9.6 minutes at the smallest grid and 291 minutes at the largest.

To benchmark SCR against an existing approach, we compared it with spCorr under the identical data-generating mechanism, across the sample sizes for which spCorr was computationally feasible ($n \leq 20{,}000$); spCorr did not scale beyond 20,000 locations, whereas SCR was applied up to 50,000. At every matched sample size, SCR recovered the spatially varying network substantially more accurately: its mean MCC (0.920, 0.935, 0.931, and 0.933 at $n = 5{,}000$, 10,000, 15,000, and 20,000) exceeded that of spCorr (0.719, 0.830, 0.821, and 0.801) by roughly 0.10 to 0.20. Figures S1–S4 provide a visual comparison between spCorr and SCR in recovering the true correlation fields. Across sample sizes, SCR consistently estimated surfaces that more faithfully followed the ground-truth fields. To supplement the foregoing visual comparisons, we summarized the field recovery performance of each method through the root-mean-squared error (RMSE), as well as the Pearson correlation between the true and estimated correlation surfaces (Figures S5–S8). Across sample sizes, SCR consistently achieved lower RMSE and higher Pearson correlation, indicating superior performance. For brevity purposes, only Replicate 1 was visualized; however, the patterns remained nearly identical across replicates. SCR also maintained markedly lower mean false positive rates (0.010–0.016 versus 0.042–0.055 for spCorr) and correspondingly higher true negative rates ($> 0.983$ versus 0.945–0.958); although spCorr attained high true positive rates for $n \geq 10{,}000$, it did so by over-selecting edges. The gap was most pronounced at the smallest grid, where spCorr's recovery degraded sharply and became highly variable (MCC 0.719, TPR 0.807, replicate standard deviation 0.338), while SCR remained stable (MCC 0.920, standard deviation 0.062). The numerical details of the foregoing comparison have been provided in Table S2. In regards to computational scalability, on the same per-edge time scale, SCR was also markedly faster (Figure S13). Its mean per-edge runtime of 1.3, 2.8, 4.7, and 7.1 seconds at $n = 5{,}000$, 10,000, 15,000, and 20,000 was roughly five to seven times faster than spCorr (8.9, 19.4, 32.8, and 37.1 seconds). Overall, SCR provided more accurate, more stable, and faster network recovery than spCorr across all sample sizes spCorr could handle, while additionally scaling to substantially larger spatial resolutions.

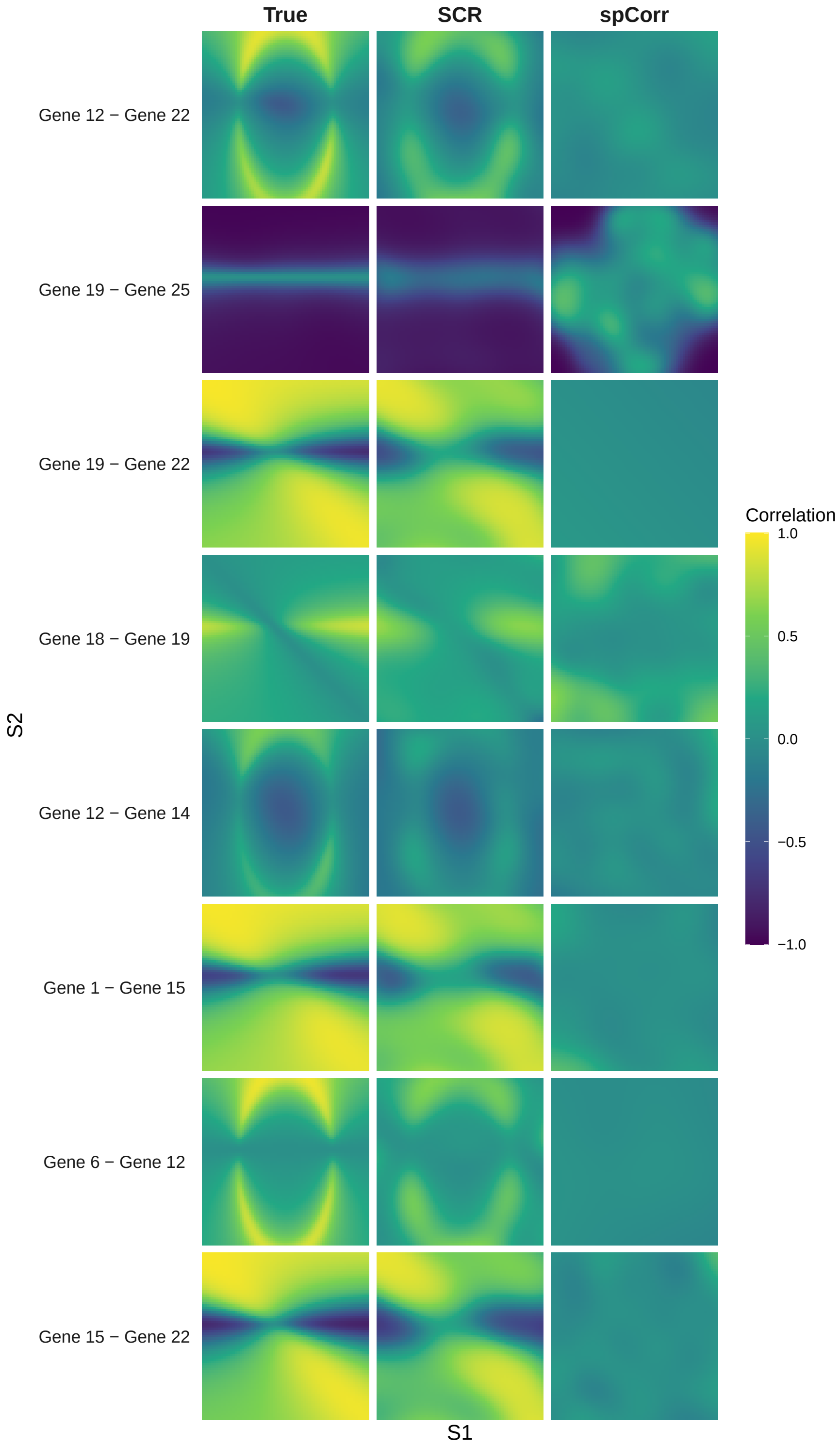

Figure S1: Each row is one edge; the three columns show the ground-truth correlation field and the fields estimated by SCR and spCorr across the $n = 5{,}000$ spatial locations of a simulated tissue. Color encodes the estimated correlation at each location (shared scale, $[-1, 1]$). Figures S1–S4 show the same quantity for varying $n$.

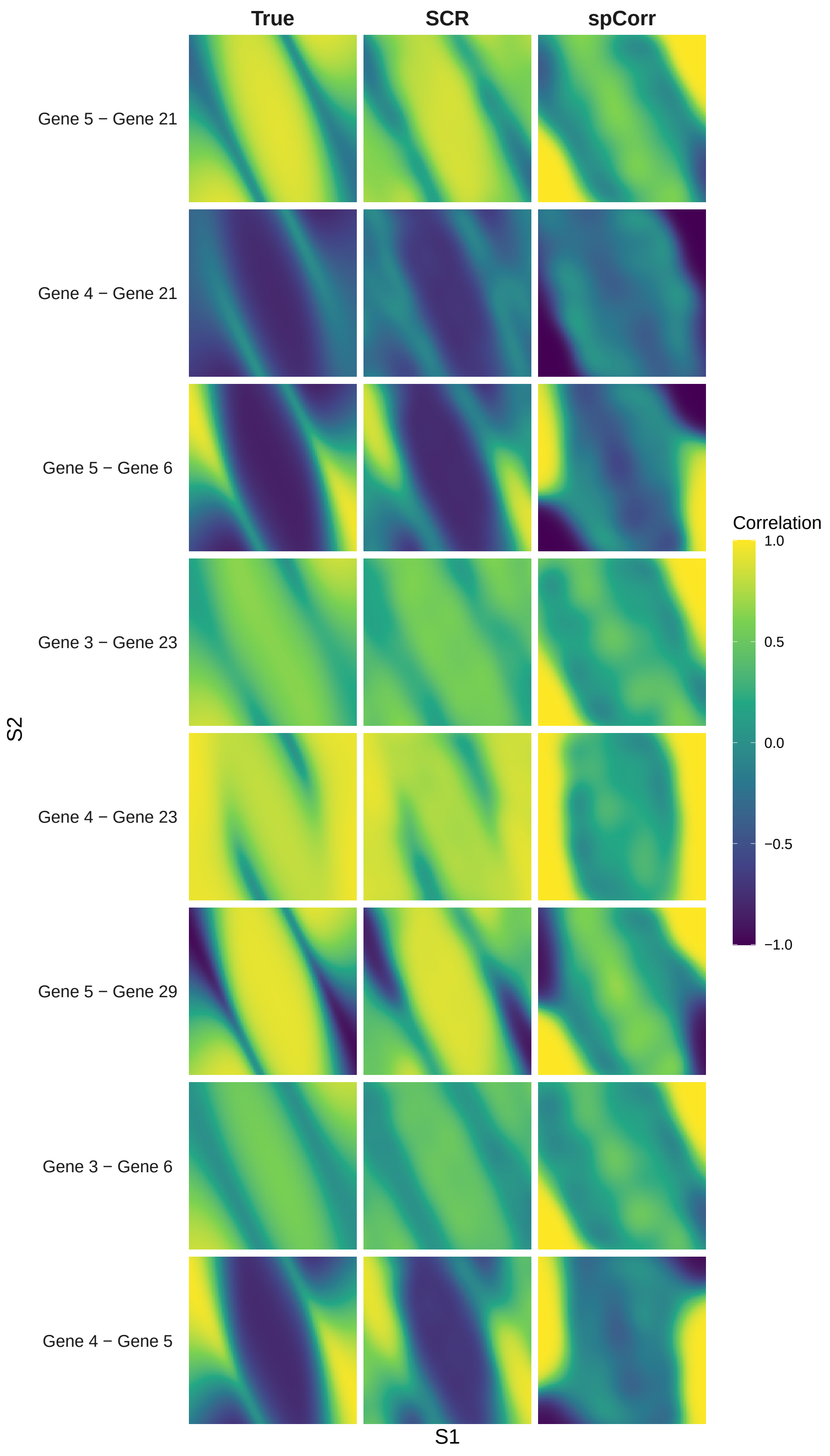

Figure S2: As in Figure S1, for $n = 10{,}000$.

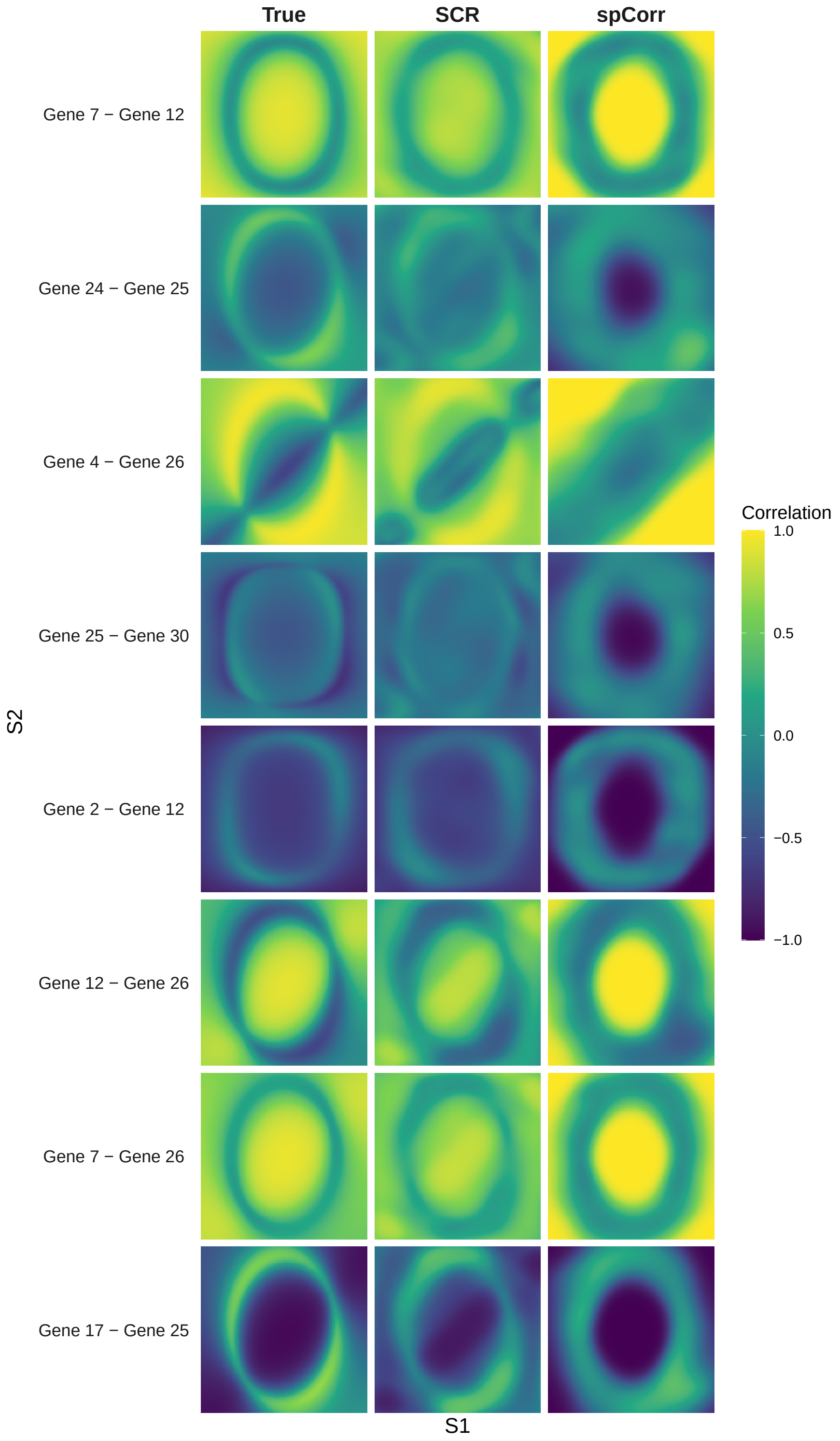

Figure S3: As in Figure S1, for $n = 15{,}000$.

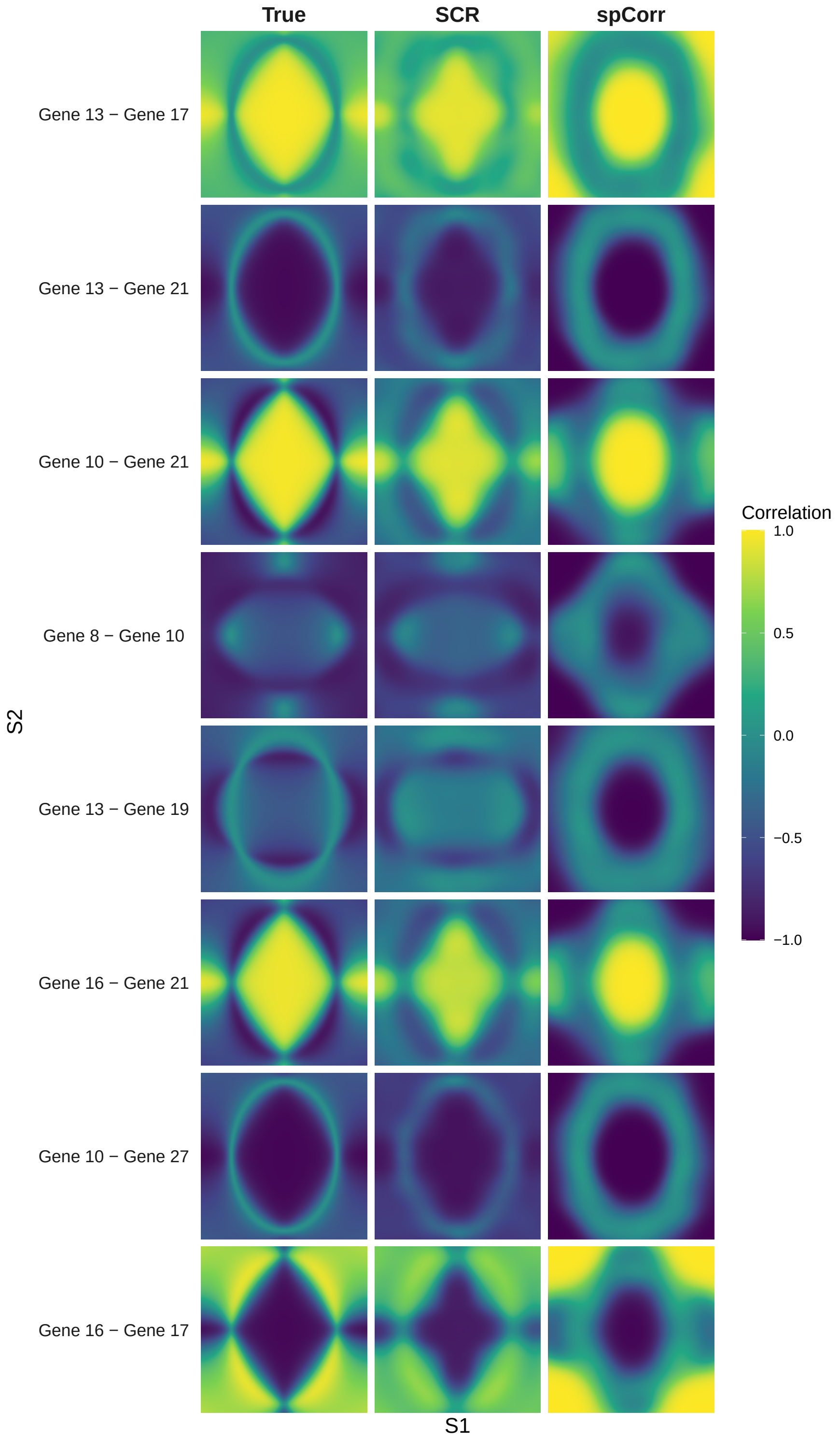

Figure S4: As in Figure S1, for $n = 20{,}000$.

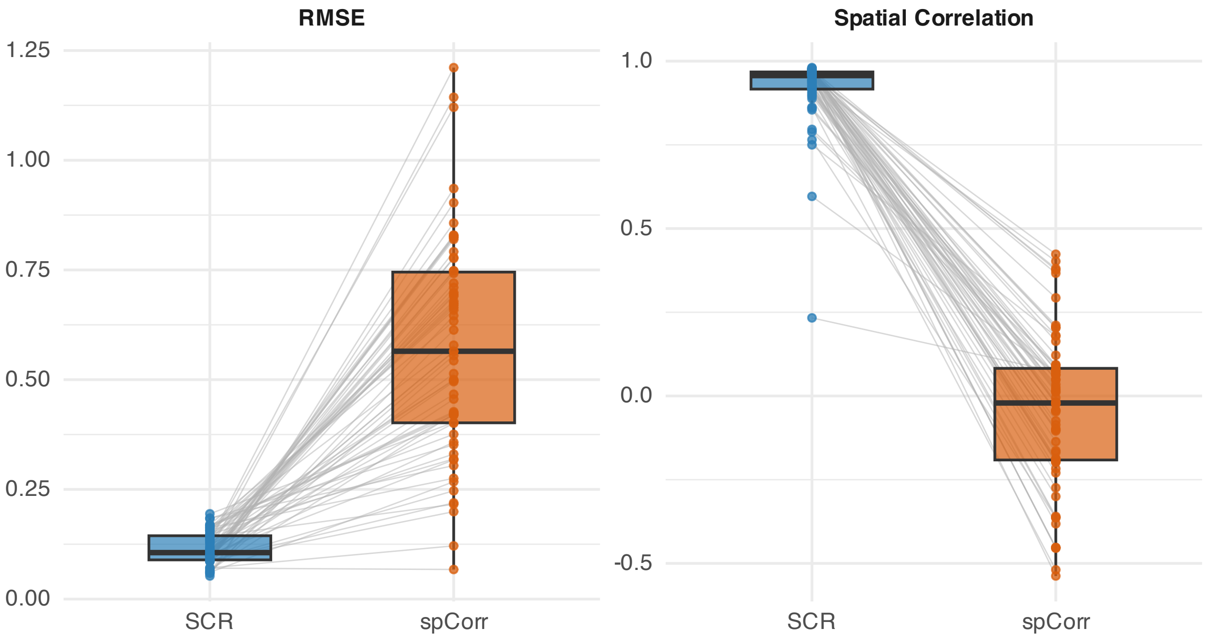

Figure S5: Each point is one SCG edge in Replicate 1 with $n = 5{,}000$, scored by comparing its estimated spatially varying correlation to the ground truth across all $n$ spatial locations; grey lines connect the same edge under the two methods (paired). *Left:* root-mean-squared error (RMSE) between the estimated and true field. *Right:* spatial correlation (Pearson) between the estimated and true field. Boxes show the median and interquartile range across edges. Figures S5–S8 show the same quantities for varying $n$.

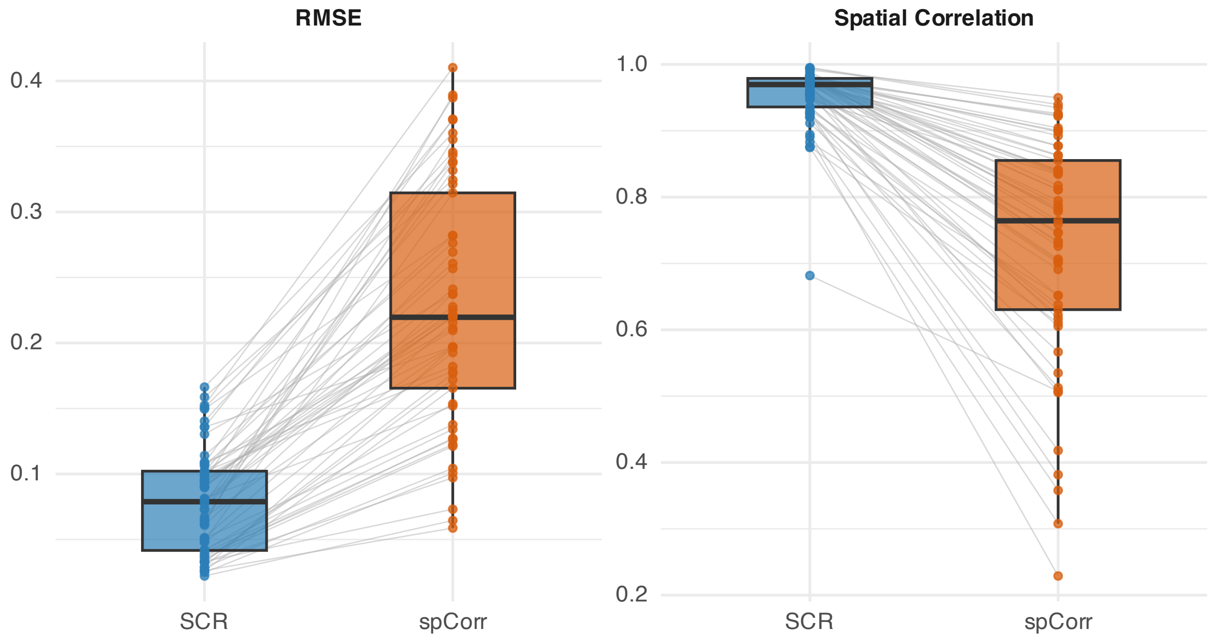

Figure S6: As in Figure S5 for $n = 10{,}000$

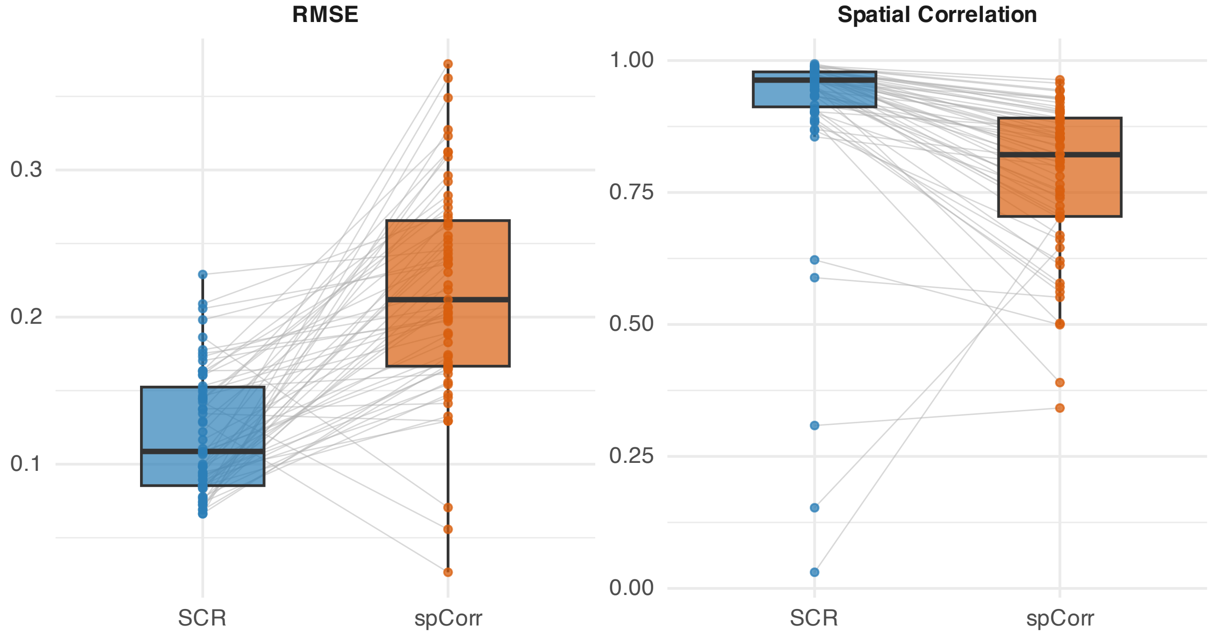

Figure S7: As in Figure S5 for $n = 15{,}000$

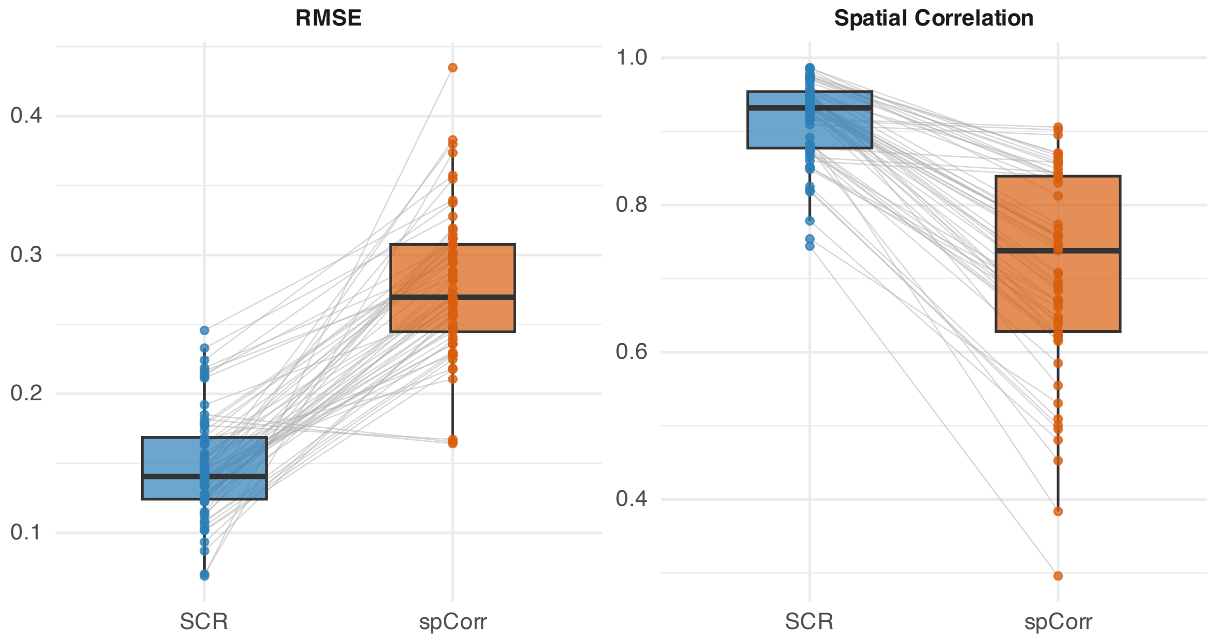

Figure S8: As in Figure S5 for $n = 20{,}000$

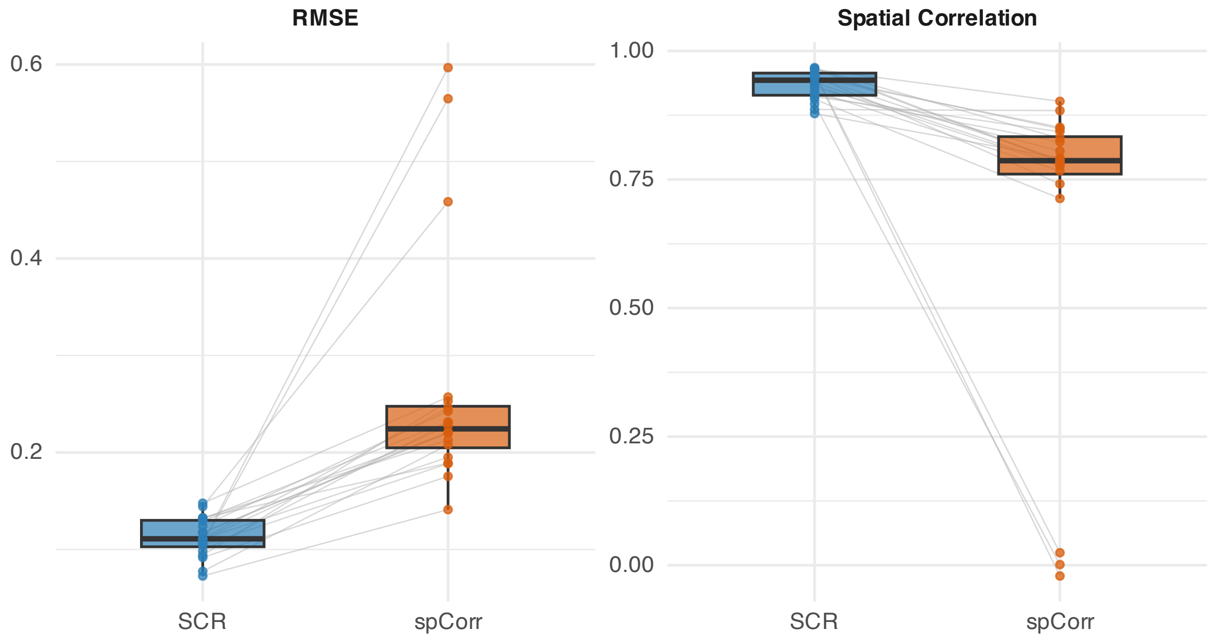

Figure S9: Each point summarizes one of the 20 replicates at $n = 5{,}000$: its median RMSE or spatial correlation, taken across that replicate's own SCG edges, comparing the estimated spatially varying correlation to the ground truth. Grey lines connect the same replicate under the two methods. *Left:* root-mean-squared error (RMSE) between the estimated and true field. *Right:* spatial correlation (Pearson) between the estimated and true field. Boxes show the median and interquartile range across replicates. Figures S9–S12 show the same quantities for varying $n$.

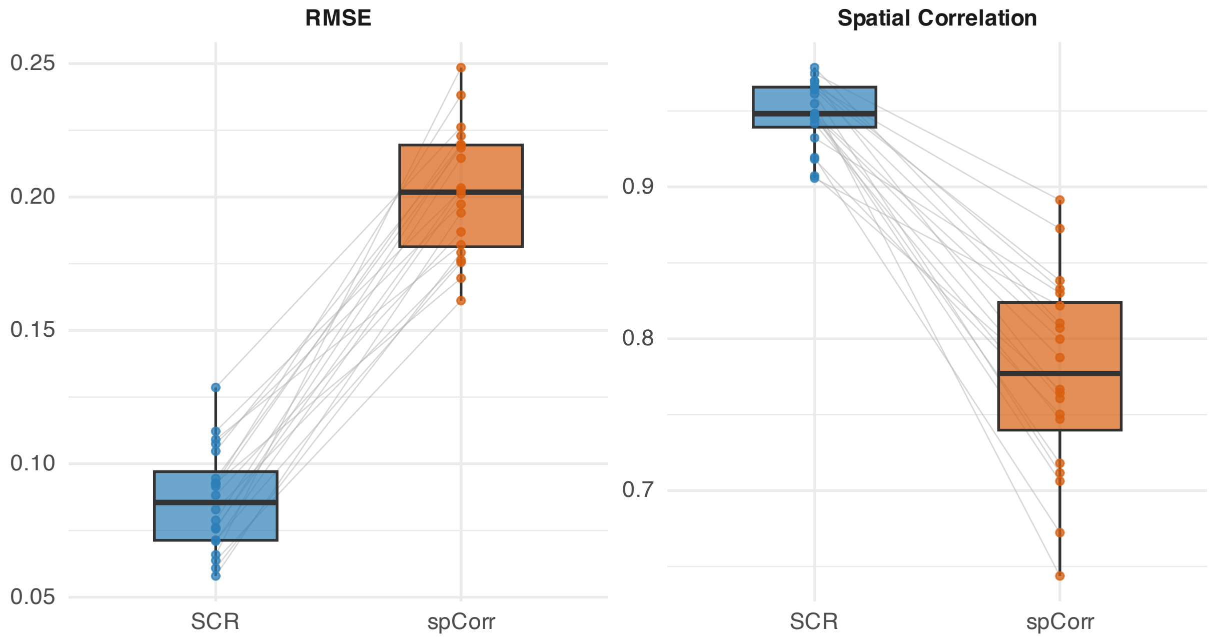

Figure S10: As in Figure S9 for $n = 10{,}000$

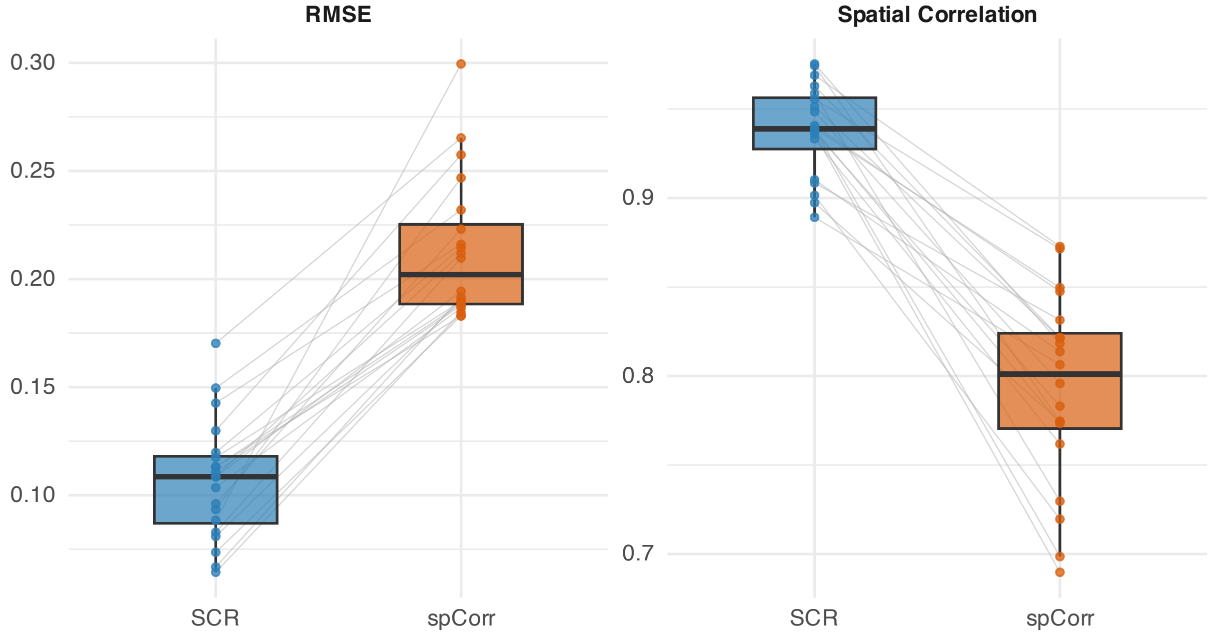

Figure S11: As in Figure S9 for $n = 15{,}000$

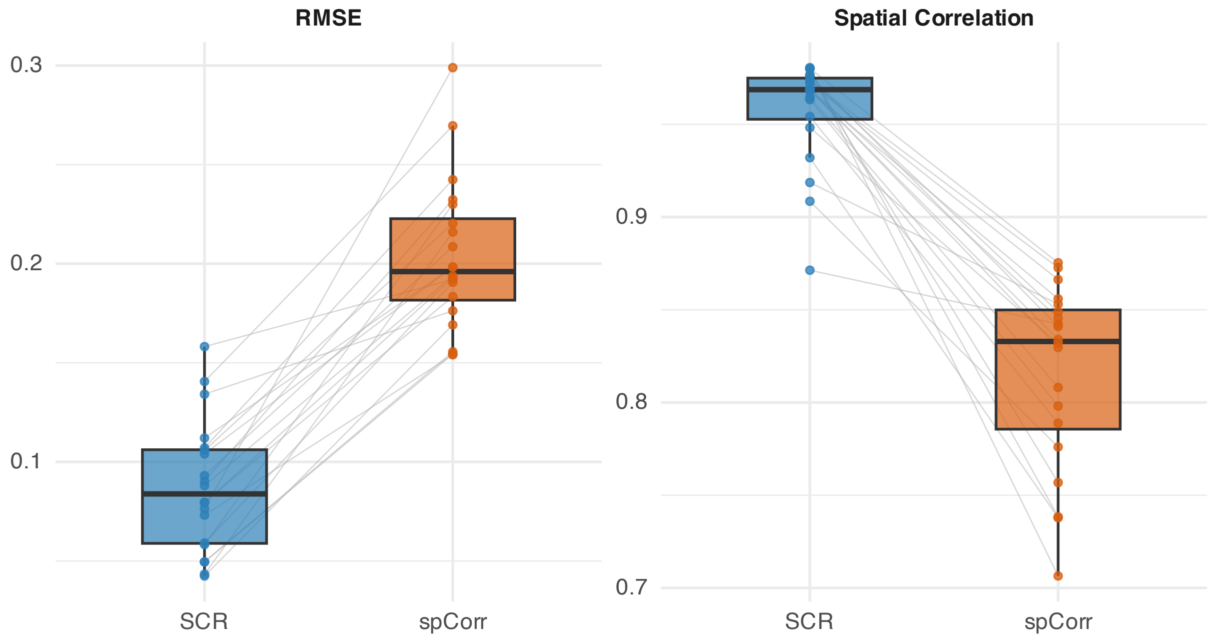

Figure S12: As in Figure S9 for $n = 20{,}000$

Table S2: Comparison of network recovery performance between SCR and spCorr across sample sizes.

| $n$ | **Method** | **TPR** | **FPR** | **MCC** | **Selection Rate** |
|---|---|---|---|---|---|
| 5,000 | SCR | 0.9804 [0.9603, 1.0000] | 0.0160 [0.0093, 0.0226] | 0.9204 [0.8933, 0.9475] | 0.1224 [0.1167, 0.1282] |
| | spCorr | 0.8066 [0.6639, 0.9493] | 0.0418 [0.0246, 0.0591] | 0.7191 [0.5709, 0.8673] | 0.1257 [0.1183, 0.1332] |
| 10,000 | SCR | 0.9723 [0.9558, 0.9888] | 0.0111 [0.0073, 0.0148] | 0.9354 [0.9211, 0.9497] | 0.1116 [0.1043, 0.1189] |
| | spCorr | 0.9872 [0.9697, 1.0000] | 0.0433 [0.0344, 0.0523] | 0.8301 [0.7991, 0.8611] | 0.1423 [0.1329, 0.1517] |
| 15,000 | SCR | 0.9759 [0.9521, 0.9997] | 0.0132 [0.0094, 0.0170] | 0.9309 [0.9125, 0.9494] | 0.1193 [0.1093, 0.1293] |
| | spCorr | 0.9976 [0.9941, 1.0000] | 0.0504 [0.0421, 0.0586] | 0.8208 [0.7967, 0.8449] | 0.1547 [0.1441, 0.1653] |
| 20,000 | SCR | 0.9741 [0.9502, 0.9980] | 0.0119 [0.0088, 0.0149] | 0.9334 [0.9183, 0.9486] | 0.1129 [0.1030, 0.1227] |
| | spCorr | 0.9984 [0.9954, 1.0000] | 0.0548 [0.0464, 0.0632] | 0.8013 [0.7747, 0.8279] | 0.1538 [0.1429, 0.1646] |

Values reported as mean [95% CI]. Abbreviations: TPR, true positive rate; FPR, false positive rate; MCC, Matthews correlation coefficient. Selection rate denotes the proportion of candidate edges selected by each method. Results are based on 20 independent replications. Confidence intervals are constructed as the mean $\pm 1.96$ standard errors and truncated to $[0, 1]$ where necessary.

Table S3: Network and correlation field recovery performance for SCR across sample sizes at the 10% FDR level.

| $n$ | **RV** | **FDR** | **TPR** | **FPR** | **MCC** | **Selection Rate** |
|---|---|---|---|---|---|---|
| 5,000 | 0.9841 [0.9821, 0.9861] | 0.1153 [0.0688, 0.1617] | 0.9804 [0.9603, 1.0000] | 0.0160 [0.0093, 0.0226] | 0.9204 [0.8933, 0.9475] | 0.1224 [0.1167, 0.1282] |
| 10,000 | 0.9898 [0.9884, 0.9912] | 0.0850 [0.0607, 0.1094] | 0.9723 [0.9558, 0.9888] | 0.0111 [0.0073, 0.0148] | 0.9354 [0.9211, 0.9497] | 0.1116 [0.1043, 0.1189] |
| 15,000 | 0.9866 [0.9841, 0.9890] | 0.0957 [0.0718, 0.1196] | 0.9759 [0.9521, 0.9997] | 0.0132 [0.0094, 0.0170] | 0.9309 [0.9125, 0.9494] | 0.1193 [0.1093, 0.1293] |
| 20,000 | 0.9900 [0.9875, 0.9926] | 0.0903 [0.0691, 0.1116] | 0.9741 [0.9502, 0.9980] | 0.0119 [0.0088, 0.0149] | 0.9334 [0.9183, 0.9486] | 0.1129 [0.1030, 0.1227] |
| 30,000 | 0.9901 [0.9868, 0.9934] | 0.0791 [0.0615, 0.0967] | 0.9617 [0.9330, 0.9904] | 0.0096 [0.0076, 0.0116] | 0.9331 [0.9204, 0.9458] | 0.1071 [0.0997, 0.1146] |
| 40,000 | 0.9888 [0.9866, 0.9911] | 0.1128 [0.0879, 0.1377] | 0.9899 [0.9823, 0.9976] | 0.0162 [0.0123, 0.0201] | 0.9284 [0.9134, 0.9434] | 0.1239 [0.1170, 0.1309] |
| 50,000 | 0.9890 [0.9869, 0.9912] | 0.0899 [0.0678, 0.1120] | 0.9890 [0.9822, 0.9958] | 0.0118 [0.0086, 0.0151] | 0.9421 [0.9282, 0.9560] | 0.1176 [0.1106, 0.1246] |

Values reported as mean [95% CI]. Abbreviations: RV, RV coefficient; FDR, false discovery rate; TPR, true positive rate; FPR, false positive rate; MCC, Matthews correlation coefficient. Selection rate denotes the proportion of candidate edges selected by SCR. Results are based on 20 independent replications at a nominal FDR level of 10%. Confidence intervals are constructed as the mean $\pm 1.96$ standard errors and truncated to $[0, 1]$ where necessary.

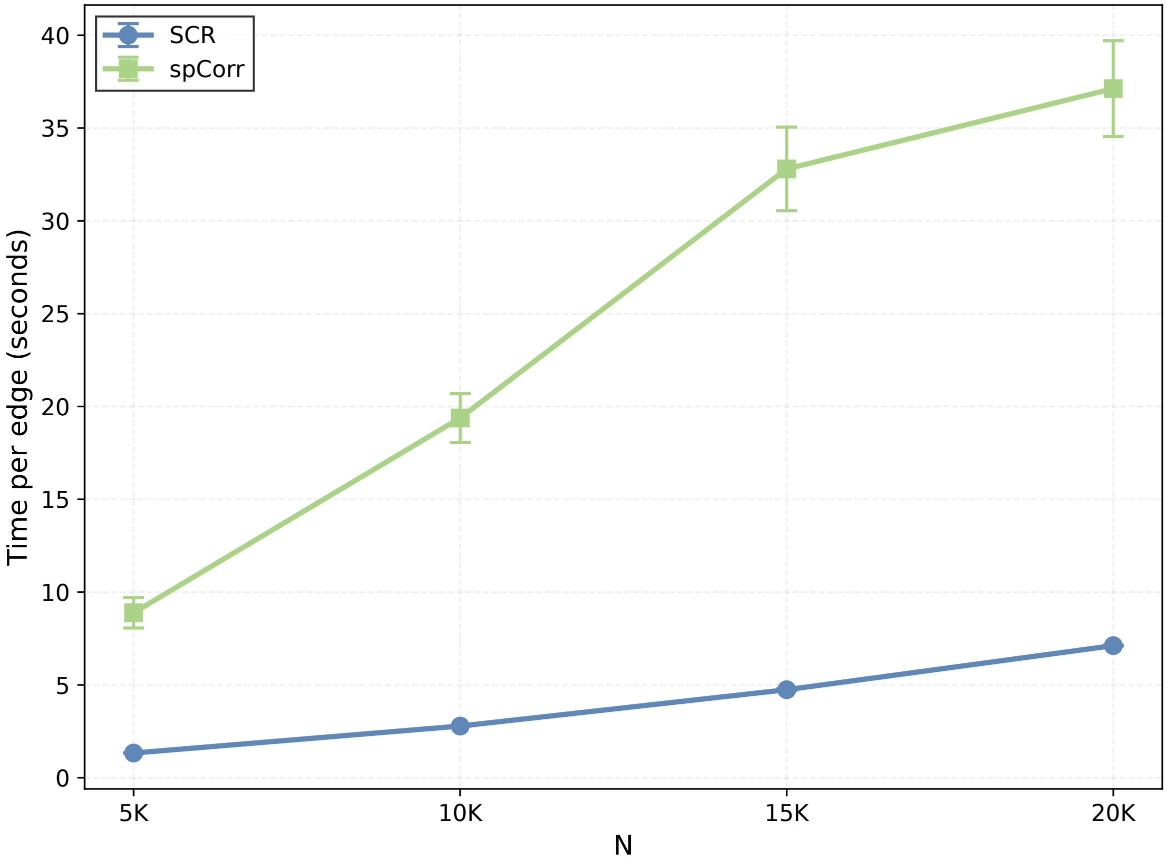

Figure S13: Computational scalability. Mean per-edge runtime (in seconds, averaged over the 435 gene pairs and 20 replicates) for SCR and spCorr as a function of the number of spatial locations $n$; error bars denote $\pm 1$ standard deviation across replicates. Error bars for SCR are not visible because its runtime was highly consistent across replicates relative to the time scale of the plot. SCR is roughly five to seven times faster than spCorr at every sample size, and spCorr was not computationally feasible beyond $n = 20{,}000$.

#### S2.1 Hardware configuration

SCR was run on an NVIDIA A100 GPU (80 GB VRAM, 4 CPUs, 128 GB RAM) on the UAB Cheaha HPC cluster; spCorr, which does not support GPU acceleration, was run using 10 CPUs with 16 GB RAM per CPU.

#### S2.2 Sensitivity analysis

We assessed the sensitivity of SCG identification to the nominal false discovery rate (FDR) threshold by repeating the analysis at target levels of 5%, 10%, and 15%, for sample sizes $n \in \{5{,}000, 10{,}000, 15{,}000, 20{,}000\}$ under the same data-generating mechanism as the primary simulation. Tables S4–S7 report the realized FDR, true positive rate (TPR), false positive rate (FPR), and Matthews correlation coefficient (MCC) for SCR and spCorr at each level. At every sample size and threshold, SCR attained substantially higher MCC and lower FPR than spCorr; because spCorr does not target a nominal FDR, it is shown at its single operating point for reference.

Table S4: Sensitivity analysis for $n = 5{,}000$

| **FDR Level** | **Method** | **FDR** | **TPR** | **FPR** | **MCC** |
|---|---|---|---|---|---|
| 5% | SCR | 0.0856 [0.0400, 0.1311] | 0.9644 [0.9402, 0.9885] | 0.0112 [0.0050, 0.0173] | 0.9292 [0.9024, 0.9560] |
| | spCorr | 0.2963 [0.1719, 0.4207] | 0.8066 [0.6639, 0.9493] | 0.0418 [0.0246, 0.0591] | 0.7191 [0.5709, 0.8673] |
| 10% | SCR | 0.1153 [0.0688, 0.1617] | 0.9804 [0.9603, 1.0000] | 0.0160 [0.0093, 0.0226] | 0.9204 [0.8933, 0.9475] |
| | spCorr | 0.2963 [0.1719, 0.4207] | 0.8066 [0.6639, 0.9493] | 0.0418 [0.0246, 0.0591] | 0.7191 [0.5709, 0.8673] |
| 15% | SCR | 0.1588 [0.1123, 0.2054] | 0.9891 [0.9729, 1.0000] | 0.0233 [0.0163, 0.0304] | 0.8986 [0.8709, 0.9263] |
| | spCorr | 0.2963 [0.1719, 0.4207] | 0.8066 [0.6639, 0.9493] | 0.0418 [0.0246, 0.0591] | 0.7191 [0.5709, 0.8673] |

Values reported as mean [95% CI]. Abbreviations: FDR, false discovery rate; TPR, true positive rate; FPR, false positive rate; MCC, Matthews correlation coefficient. spCorr does not target a nominal FDR level, so its single operating point is repeated at each row for reference. Results based on 20 independent replications.

Table S5: Sensitivity analysis for $n = 10{,}000$

| FDR Level | Method | FDR | TPR | FPR | MCC |
|---|---|---|---|---|---|
| 5% | SCR | 0.0508 [0.0294, 0.0721] | 0.9548 [0.9332, 0.9765] | 0.0063 [0.0033, 0.0093] | 0.9455 [0.9311, 0.9600] |
| | spCorr | 0.2657 [0.2206, 0.3108] | 0.9872 [0.9697, 1.0000] | 0.0433 [0.0344, 0.0523] | 0.8301 [0.7991, 0.8611] |
| 10% | SCR | 0.0850 [0.0607, 0.1094] | 0.9723 [0.9558, 0.9888] | 0.0111 [0.0073, 0.0148] | 0.9354 [0.9211, 0.9497] |
| | spCorr | 0.2657 [0.2206, 0.3108] | 0.9872 [0.9697, 1.0000] | 0.0433 [0.0344, 0.0523] | 0.8301 [0.7991, 0.8611] |
| 15% | SCR | 0.1234 [0.0972, 0.1496] | 0.9857 [0.9750, 0.9964] | 0.0167 [0.0125, 0.0209] | 0.9201 [0.9060, 0.9343] |
| | spCorr | 0.2657 [0.2206, 0.3108] | 0.9872 [0.9697, 1.0000] | 0.0433 [0.0344, 0.0523] | 0.8301 [0.7991, 0.8611] |

Values reported as mean [95% CI]. Abbreviations: FDR, false discovery rate; TPR, true positive rate; FPR, false positive rate; MCC, Matthews correlation coefficient. spCorr does not target a nominal FDR level, so its single operating point is repeated at each row for reference. Results based on 20 independent replications.

Table S6: Sensitivity analysis for $n = 15{,}000$

| FDR Level | Method | FDR | TPR | FPR | MCC |
|---|---|---|---|---|---|
| 5% | SCR | 0.0606 [0.0406, 0.0807] | 0.9645 [0.9386, 0.9905] | 0.0079 [0.0050, 0.0107] | 0.9452 [0.9277, 0.9627] |
| | spCorr | 0.2867 [0.2506, 0.3229] | 0.9976 [0.9941, 1.0000] | 0.0504 [0.0421, 0.0586] | 0.8208 [0.7967, 0.8449] |
| 10% | SCR | 0.0957 [0.0718, 0.1196] | 0.9759 [0.9521, 0.9997] | 0.0132 [0.0094, 0.0170] | 0.9309 [0.9125, 0.9494] |
| | spCorr | 0.2867 [0.2506, 0.3229] | 0.9976 [0.9941, 1.0000] | 0.0504 [0.0421, 0.0586] | 0.8208 [0.7967, 0.8449] |
| 15% | SCR | 0.1403 [0.1154, 0.1653] | 0.9852 [0.9650, 1.0000] | 0.0205 [0.0160, 0.0250] | 0.9092 [0.8913, 0.9271] |
| | spCorr | 0.2867 [0.2506, 0.3229] | 0.9976 [0.9941, 1.0000] | 0.0504 [0.0421, 0.0586] | 0.8208 [0.7967, 0.8449] |

Values reported as mean [95% CI]. Abbreviations: FDR, false discovery rate; TPR, true positive rate; FPR, false positive rate; MCC, Matthews correlation coefficient. spCorr does not target a nominal FDR level, so its single operating point is repeated at each row for reference. Results based on 20 independent replications.

### S3 Real Data Analysis: Breast Cancer

#### S3.1 Data preprocessing

We compared the behavior of the ten cancer hallmark pathways together with the epithelial–mesenchymal transition (EMT) pathway in the HER2-amplified invasive ductal carcinoma sample. Genes with zero counts across all spots, and spots with zero total counts, were removed. Counts were then normalized using the variance-stabilizing transformation implemented in `sctransform` Hafemeister and Satija (2019), run with `vst.flavor = "v2"` and `n_genes = NULL` so that model parameters were estimated from all genes rather than from a subsample. Briefly, this approach fits a regularized negative binomial regression per gene with the log total UMI count per spot as covariate and returns Pearson residuals.

Downstream analysis was restricted to genes shared between the normalized matrix and the pathway gene sets. Namely, the genes implicated in the cancer hallmarks collection Menyhárt et al. (2025), and those implicated in the hallmark EMT set Liberzon et al. (2015). Within each set, genes detected in fewer than 1% of spots in the raw counts were further excluded. Resulting gene counts per pathway are given in Supplementary Table S8.

#### S3.2 Section overview

Table S9 summarizes the number of total possible edges, computed as $\binom{p}{2}$, as well as the number of edges identified as SCG and the corresponding proportion across the pathways examined for the breast cancer data application. As expected, SCGs constituted a small proportion of the total possible edges, ranging from approximately 2% to 20%.

Table S10 presents the region-stratified network connectivity scores (CS). Briefly, CS is computed as the number of edges in a given network that exceed 0.1 in magnitude, normalized by the maximum count across spots and pathways to yield a score that ranges from 0, indicating no connectivity, to 1, indicating high relative connectivity. Connectivity in this context is taken to represent pathway-level coordinated activity, with high values indicating strong coordination. Examining the CS scores across pathways and regions of the tumor sample, we note that many pathways associated with tumorigenesis, such as tissue invasion and metastasis, sustaining proliferative signaling, and epithelial-mesenchymal transition, are highly connected in the tumor region. Certain pathways, such as evading immune destruction, sustained angiogenesis, and tumor-promoting inflammation, appear most connected in the intermediate regions. Reprogramming energy metabolism was the only pathway that had higher average connectivity in normal regions than in both intermediate and tumor regions. Figure S17 visualizes CS in a spatially varying manner stratified by pathway. Broadly, patterns similar to those in Table S10 are observed, with most pathways showing high connectivity in tumor regions.

Figures S14–S16 visualise the top five edges by adjusted Rand index (ARI) for each of the pathways examined. In each cell, the top surface corresponds to the estimated correlation field between two genes across the breast cancer sample, with height corresponding to the magnitude and color corresponding to the direction of correlation. Underlying each correlation field is the corresponding sample with tissue annotations. Notable patterns emerge when the correlation surfaces are interpreted alongside the underlying tissue annotations.

Table S7: Sensitivity analysis for $n = 20{,}000$

| **FDR Level** | **Method** | **FDR** | **TPR** | **FPR** | **MCC** |
|---|---|---|---|---|---|
| 5% | SCR | 0.0545 [0.0387, 0.0702] | 0.9596 [0.9333, 0.9860] | 0.0067 [0.0048, 0.0086] | 0.9463 [0.9320, 0.9605] |
| | spCorr | 0.3173 [0.2786, 0.3560] | 0.9984 [0.9954, 1.0000] | 0.0548 [0.0464, 0.0632] | 0.8013 [0.7747, 0.8279] |
| 10% | SCR | 0.0903 [0.0691, 0.1116] | 0.9741 [0.9502, 0.9980] | 0.0119 [0.0088, 0.0149] | 0.9334 [0.9183, 0.9486] |
| | spCorr | 0.3173 [0.2786, 0.3560] | 0.9984 [0.9954, 1.0000] | 0.0548 [0.0464, 0.0632] | 0.8013 [0.7747, 0.8279] |
| 15% | SCR | 0.1303 [0.1070, 0.1536] | 0.9840 [0.9663, 1.0000] | 0.0179 [0.0143, 0.0215] | 0.9151 [0.9021, 0.9281] |
| | spCorr | 0.3173 [0.2786, 0.3560] | 0.9984 [0.9954, 1.0000] | 0.0548 [0.0464, 0.0632] | 0.8013 [0.7747, 0.8279] |

Values reported as mean [95% CI]. Abbreviations: FDR, false discovery rate; TPR, true positive rate; FPR, false positive rate; MCC, Matthews correlation coefficient. spCorr does not target a nominal FDR level, so its single operating point is repeated at each row for reference. Results based on 20 independent replications.

Table S8: Pathway-level gene counts in the breast cancer sample. Total and retained gene counts are reported for each hallmark pathway.

| Pathway | Total Genes | Overlap with Dataset |
|---|---|---|
| Tissue invasion and metastasis | 624 | 428 |
| Reprogramming energy metabolism | 155 | 131 |
| Replicative immortality | 94 | 78 |
| Genome instability | 178 | 150 |
| Resisting cell death | 423 | 308 |
| Evading growth suppressors | 407 | 287 |
| Evading immune destruction | 106 | 82 |
| Sustained angiogenesis | 117 | 81 |
| Sustaining proliferative signaling | 655 | 433 |
| Tumor-promoting inflammation | 66 | 51 |
| Epithelial–mesenchymal transition | 200 | 152 |

Total genes: number of genes reported as part of the cancer hallmarks collection Menyhárt et al. (2025), and of the hallmark EMT set Liberzon et al. (2015). Overlap with dataset: genes remaining after the preprocessing steps outlined in Supplementary Section S3.1.

Table S9: Pathway-level edge selection summary in the breast cancer sample. Total and selected gene-pair edges are reported for each hallmark pathway, along with the proportion of edges selected.

| Pathway | Total Edges | Selected Edges | Proportion |
|---|---|---|---|
| Tissue invasion and metastasis | 91,378 | 6,898 | 0.0755 |
| Reprogramming energy metabolism | 8,515 | 186 | 0.0218 |
| Replicative immortality | 3,003 | 130 | 0.0433 |
| Genome instability | 11,175 | 229 | 0.0205 |
| Resisting cell death | 47,278 | 2,979 | 0.0630 |
| Evading growth suppressors | 41,041 | 2,946 | 0.0718 |
| Evading immune destruction | 3,321 | 367 | 0.1105 |
| Sustained angiogenesis | 3,240 | 663 | 0.2046 |
| Sustaining proliferative signaling | 93,528 | 6,555 | 0.0701 |
| Tumor-promoting inflammation | 1,275 | 120 | 0.0941 |
| Epithelial–mesenchymal transition | 11,476 | 2,523 | 0.2199 |

Total edges: all gene-pair edges within the pathway. Selected edges: edges identified as spatially co-expressed at the nominal FDR threshold.

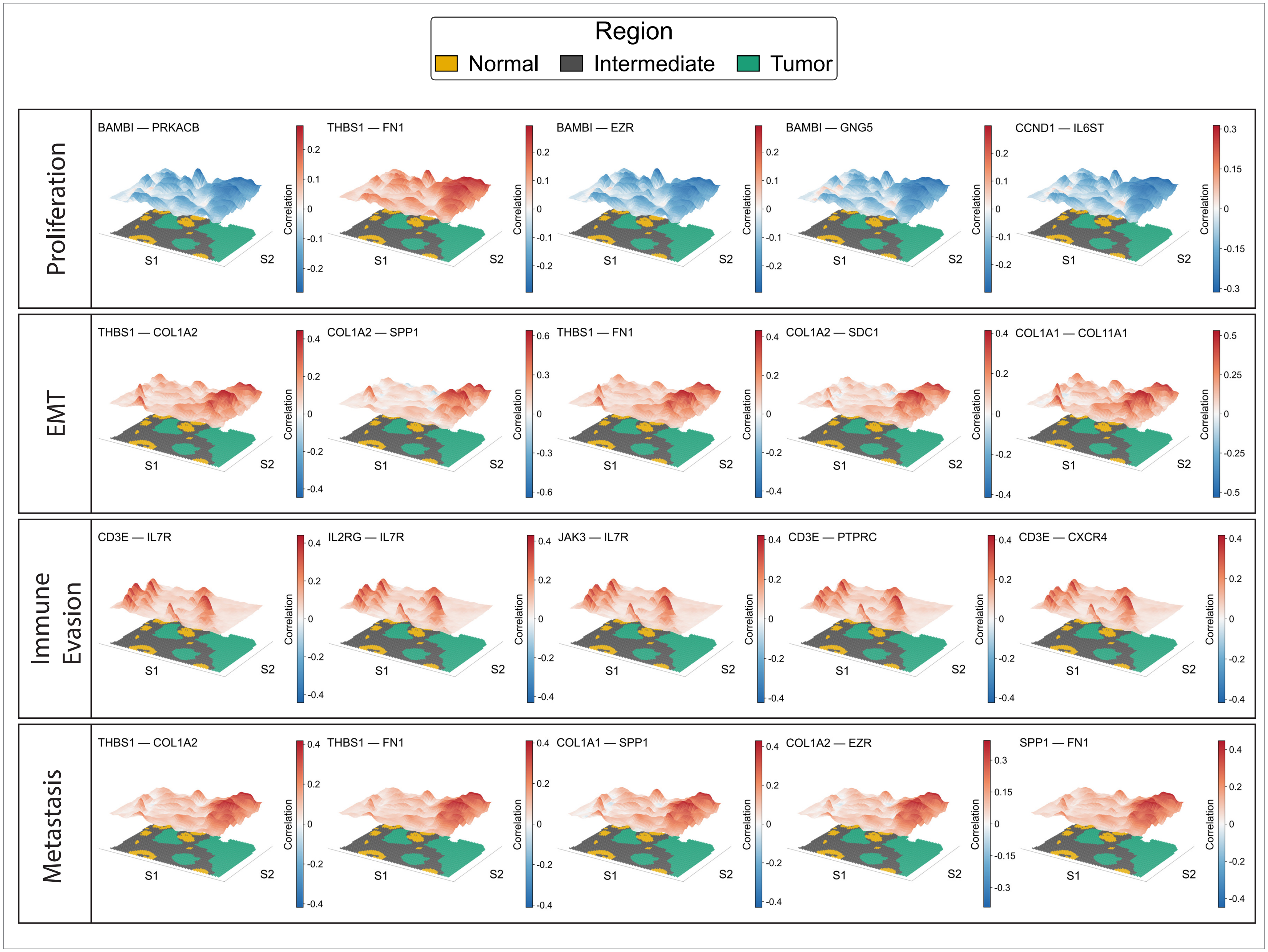

Figure S14: Spatially varying co-expression edges in the breast cancer sample. Each panel maps the estimated posterior mean correlation of one gene pair across spatial locations, with red and blue denoting positive and negative co-expression, respectively. Within each hallmark pathway, the gene-pair edges shown are those with the highest scaled adjusted Rand index (sARI) with respect to the pathologist annotations. Rows group gene pairs by pathway (proliferation, epithelial–mesenchymal transition, immune evasion, and metastasis); annotated normal, intermediate, and tumor regions are outlined for reference. Figures S14–S16 show the same sample, split across figures by pathway group.

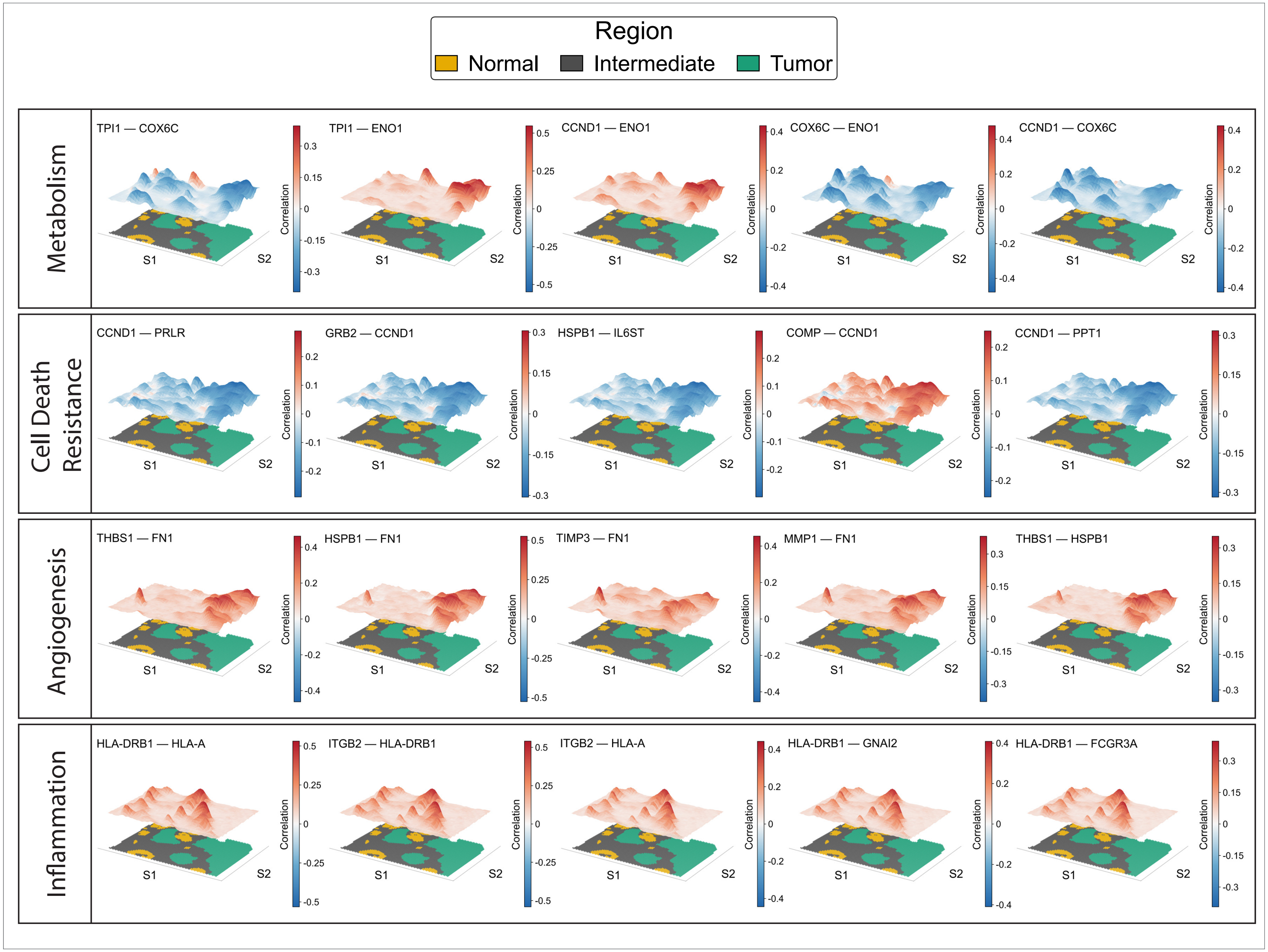

Figure S15: As in Figure S14, for the same breast cancer sample: the estimated posterior mean correlation of gene pairs is mapped across the spatial locations. Rows group gene pairs by hallmark pathway (metabolism, cell death resistance, angiogenesis, and inflammation); within each pathway, the edges shown are those with the highest sARI.

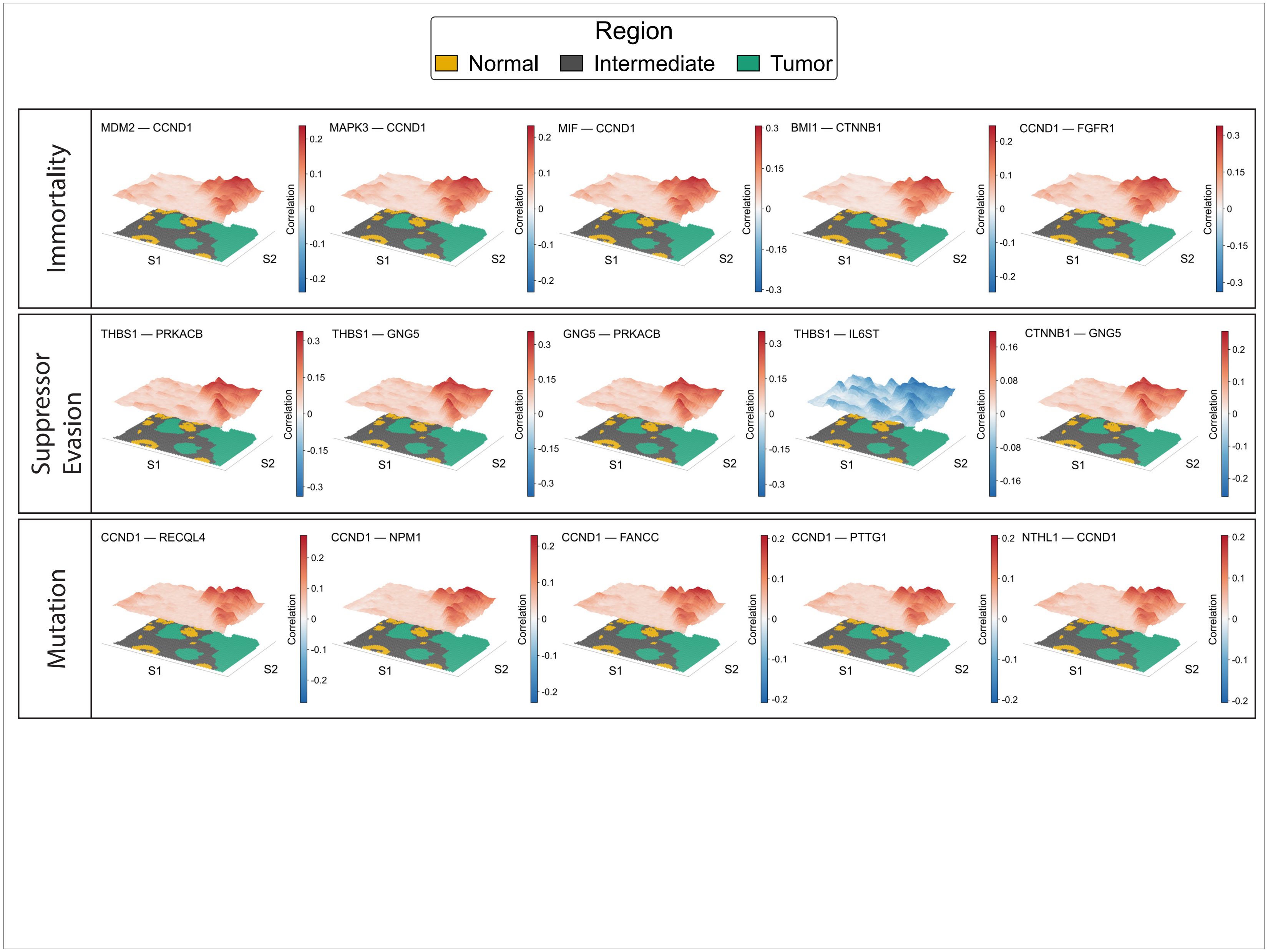

Figure S16: As in Figure S14, for the same breast cancer sample, showing the remaining hallmark pathways. Rows group gene pairs by pathway (replicative immortality, evading growth suppressors, and genome instability); within each pathway, the edges shown are those with the highest sARI.

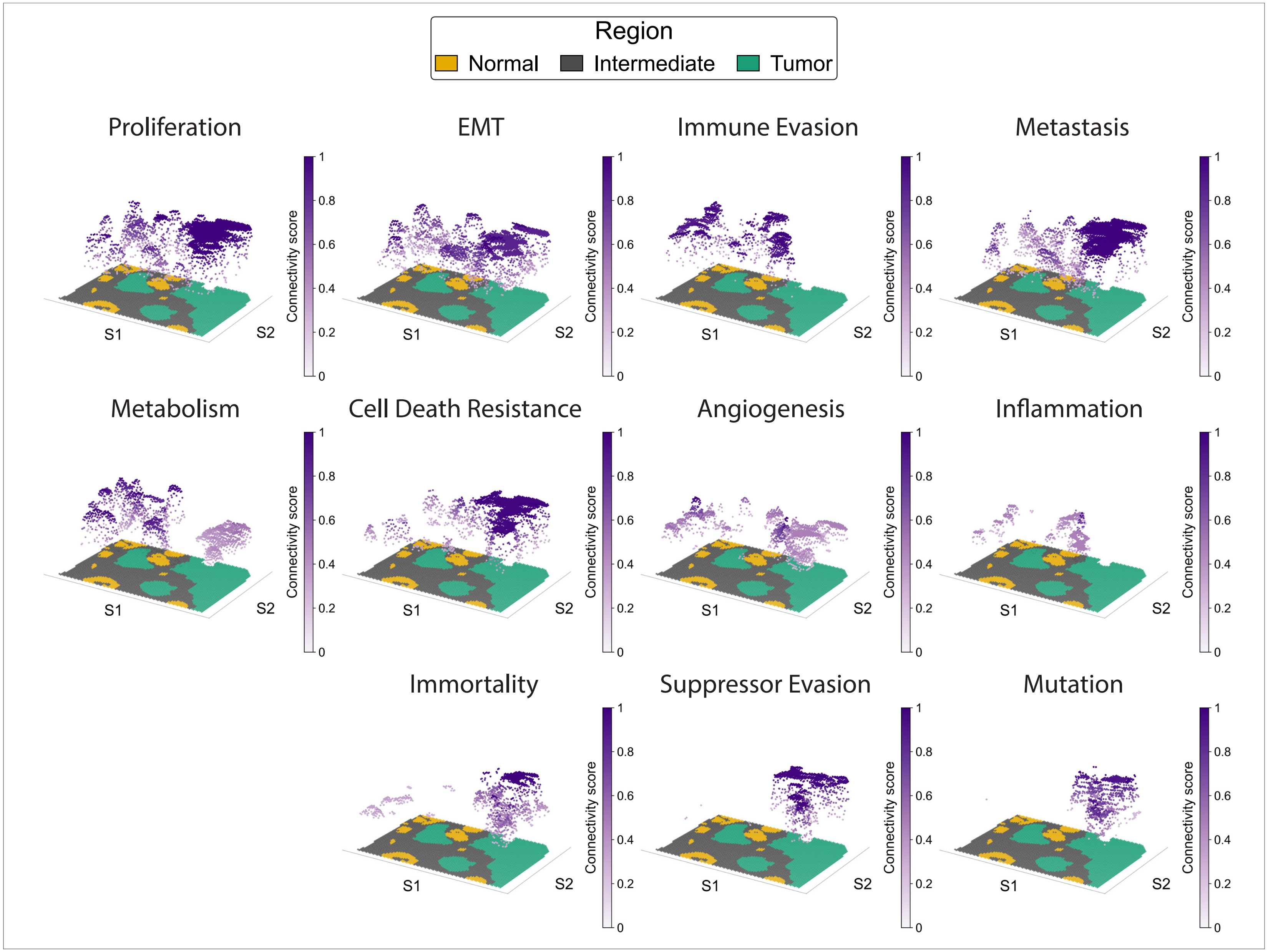

Figure S17: Spatially varying network connectivity by pathway in the breast cancer sample. Each panel shows the scaled connectivity score (CS) for a hallmark pathway across the tissue, defined as the number of edges present at a spot divided by the maximum number of edges observed across all pathways and spots. Higher CS are shown in purple, and lower CS in white. Pathologist-annotated normal, intermediate, and tumor regions are outlined. Spots with CS $< 0.1$ are not visualized no for clarity.

Table S10: Region-stratified network connectivity by hallmark pathway in the breast cancer sample. Each cell reports the mean scaled connectivity score across spots in the indicated region, with the standard deviation in parentheses; for each pathway, the region with the highest mean connectivity is shown in bold.

| **Pathway** | **Normal** | **Intermediate** | **Tumor** |
|---|---|---|---|
| Tissue invasion and metastasis | 0.36 (0.23) | 0.20 (0.23) | **0.73 (0.34)** |
| Reprogramming energy metabolism | **0.62 (0.27)** | 0.16 (0.20) | 0.35 (0.22) |
| Replicative immortality | 0.03 (0.02) | 0.08 (0.06) | **0.29 (0.17)** |
| Genome instability | 0.02 (0.00) | 0.02 (0.01) | **0.13 (0.07)** |
| Resisting cell death | 0.33 (0.23) | 0.09 (0.13) | **0.68 (0.37)** |
| Evading growth suppressors | 0.08 (0.06) | 0.06 (0.06) | **0.51 (0.39)** |
| Evading immune destruction | 0.38 (0.36) | **0.70 (0.34)** | 0.24 (0.26) |
| Sustained angiogenesis | 0.11 (0.13) | **0.31 (0.24)** | 0.22 (0.15) |
| Sustaining proliferative signaling | 0.58 (0.32) | 0.12 (0.19) | **0.75 (0.32)** |
| Tumor-promoting inflammation | 0.10 (0.09) | **0.22 (0.18)** | 0.07 (0.07) |
| Epithelial–mesenchymal transition | 0.42 (0.30) | 0.29 (0.27) | **0.59 (0.29)** |

### S4 Real Data Analysis: Alzheimer's Disease

#### S4.1 Data preprocessing

The second application uses two mouse brain sections profiled with the 10x Genomics Xenium platform. Negative control probes and blank codewords were removed from the gene set. Thereafter, cells with fewer than 50 detected transcripts were excluded, which typically corresponded to cells that were disconnected from the main tissue sample. Counts were then normalized using the variance-stabilizing transformation implemented in `sctransform`, run with `vst.flavor = "v2"` and `n_genes = NULL` so that model parameters were estimated from all genes rather than from a subsample Hafemeister and Satija (2019). Briefly, this approach fits a regularized negative binomial regression per gene with the log total transcript count per cell as covariate and returns Pearson residuals. Downstream analysis was restricted to genes in the PIG pathway gene set that were common between the wild-type and mutant samples Chen et al. (2020).

Cell-level labels were transferred from `well_11` of the STARmap PLUS mouse Alzheimer's disease atlas, using the `Main_molecular_cell_type` assignments Zeng et al. (2023). For each sample, coordinates were first brought into approximate correspondence with the reference by rescaling and reflecting the coordinates, after which each section was aligned to the reference independently using STalign Clifton et al. (2023). Briefly, STalign rasterizes point positions into continuous images and estimates an affine transformation followed by a large deformation diffeomorphic metric mapping, accommodating the non-linear distortions required to map one tissue into the coordinate space of the other. Reference labels were transferred to the aligned coordinates and subsequently refined by 5-nearest-neighbor classification with weights inversely proportional to Euclidean distance as implemented in `scikit-learn` Pedregosa et al. (2011). Label alignment prior to aggregation is presented in Supplementary Figure S18. Regions were aggregated to coarse anatomical classes using the hierarchical Allen Mouse Brain ontology, collapsing subdivisions that share a common higher-level parent (for example, cerebellar layers to "Cerebellum," olfactory bulb layers to "Olfactory bulb," and dentate gyrus plus CA fields to "Hippocampus") Wang et al. (2020). Labels spanning multiple higher-level structures were retained as composite classes. The resulting labels are shown in Supplementary Table S11.

#### S4.2 Section overview

Figure S18 depicts the two samples that were examined in the study on either side of the reference brain, which was used to label the samples we examined. Each color denotes a certain anatomical label. For downstream analyses, these region-level annotations were collapsed to appropriate parent-level regions for simpler interpretation. The correspondence between the original labels obtained from the reference brain and their mapping to the collapsed labels is provided in Table S11. Figures S19–S20 depict the 20 SCGs with the highest sARI for the MUT and WT samples, respectively. In each cell, the top surface corresponds to the estimated correlation field, with intensity encoding magnitude and color encoding direction. Underlying each correlation field is the corresponding sample with region annotations. Broadly, the MUT sample shows strong positive correlations among edges that are stronger in cortical regions compared to subcortical regions, with very few edges showing negative correlations across the brain. Among the WT figures, the cortical-to-subcortical gradient noted among MUT edges is not as prominent. Most edges show strong, localized, positive correlations. Similarly, a small number of edges show negative correlations across the tissue domain. Similar to Figures S19–S20, Figures S23–S24 depict the estimated edges for a sample of gene pairs that were identified as being co-expressed by Chen et al. (2020).

Our findings build on those of Chen et al. (2020) by resolving the co-expression pattern at spot resolution and providing a means to visualize these patterns for biological interpretation.

Figures S21–S22 compare the connectivity patterns for a given gene across the MUT and WT samples. The connectivity score (CS) computed and visualized here is complementary to that presented in Figure S17, which depicts network-level connectivity. Here, we depict gene-level connectivity, which is computed as the number of edges incident to a vertex (i.e., gene) that exceed 0.1 in magnitude, normalized by the maximum connectivity observed across samples, genes, and spots. This yields a score between 0, indicating low connectivity, and 1, indicating high relative connectivity. Each cell contains three surfaces: the top one is the CS for MUT, the middle is the CS for WT, and the bottom is the labeled brain for the WT sample. Broadly, across the genes, we see stronger connectivity in the MUT sample compared to WT. This is plausible given how the activation of the genes in the PIG pathway is associated with plaque burden.

Table S11: Mapping of reference labels to coarse anatomical regions. Each label distributed with the reference dataset is listed alongside the region to which it was assigned.

| **Original label** | **Merged region** |
|---|---|
| CTX_1 | Cortex |
| LSX_HY_MB_HB | LSX/HY/MB/HB |
| CBX_1 | Cerebellum |
| STR | Striatum |
| OB_1 | Olfactory bulb |
| FbTrt | Fiber tracts |
| L1_HPFmo_Mngs | HPF/Meninges |
| CTX_2 | Cortex |
| OB_2 | Olfactory bulb |
| TH | Thalamus |
| CBX_2 | Cerebellum |
| Hbl_VS | Hbl/VS |
| DG | Hippocampus |
| HPF_CA | Hippocampus |
| HY | Hypothalamus |
| MYdp | Brainstem |
| ENTm | Cortex |

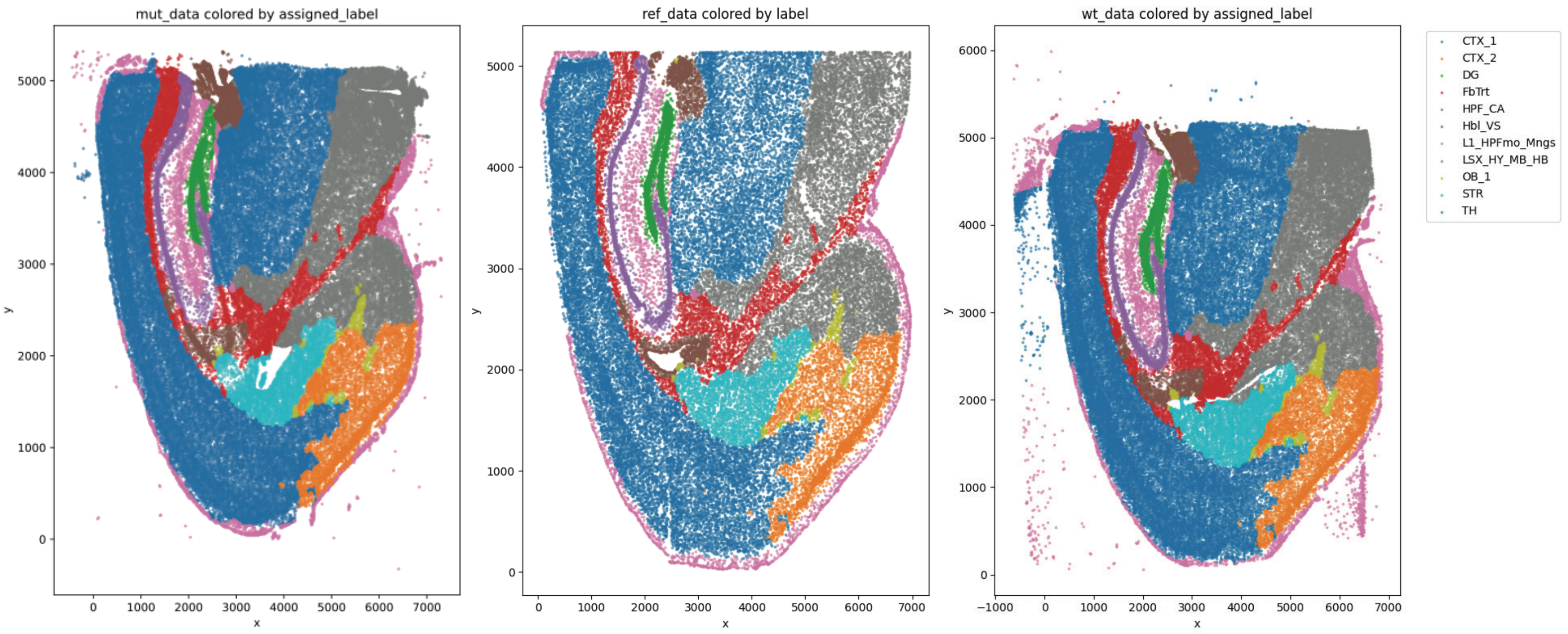

Figure S18: Anatomical label transfer for the Alzheimer's disease mouse-model brain sections. The mutant (MUT, left) and wild-type (WT, right) sections are shown after registration to the reference (middle), with each cell colored by its assigned anatomical label. Labels are shown prior to aggregation into coarse regions.

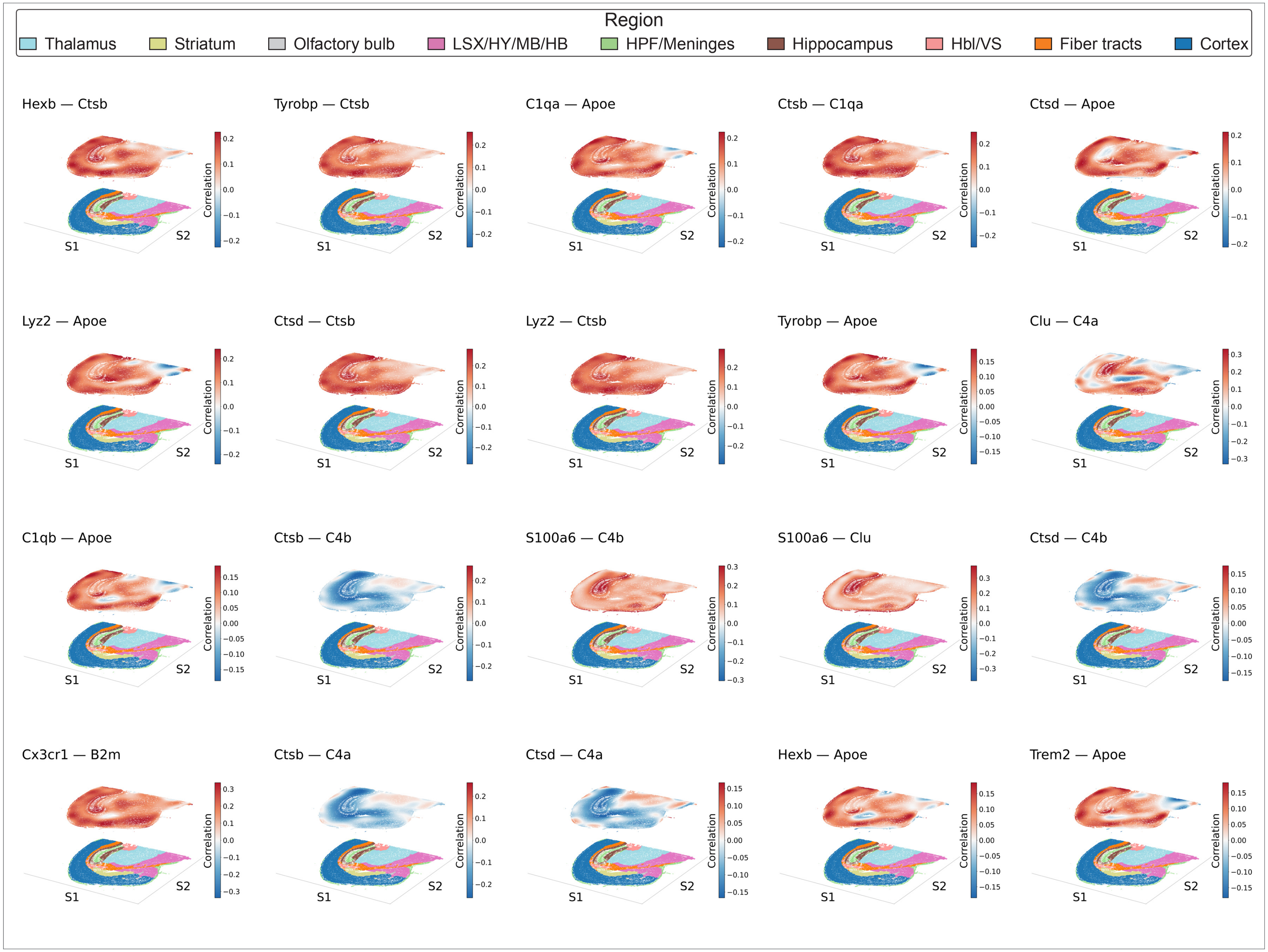

Figure S19: Spatially varying co-expression edges for selected gene pairs in the Alzheimer's disease mouse-model brain (mutant, MUT). Each panel maps the estimated posterior mean correlation of one gene pair across a coronal section, with red and blue denoting positive and negative co-expression. Anatomical regions are indicated in the legend. The 20 edges shown are those with the highest sARI values.

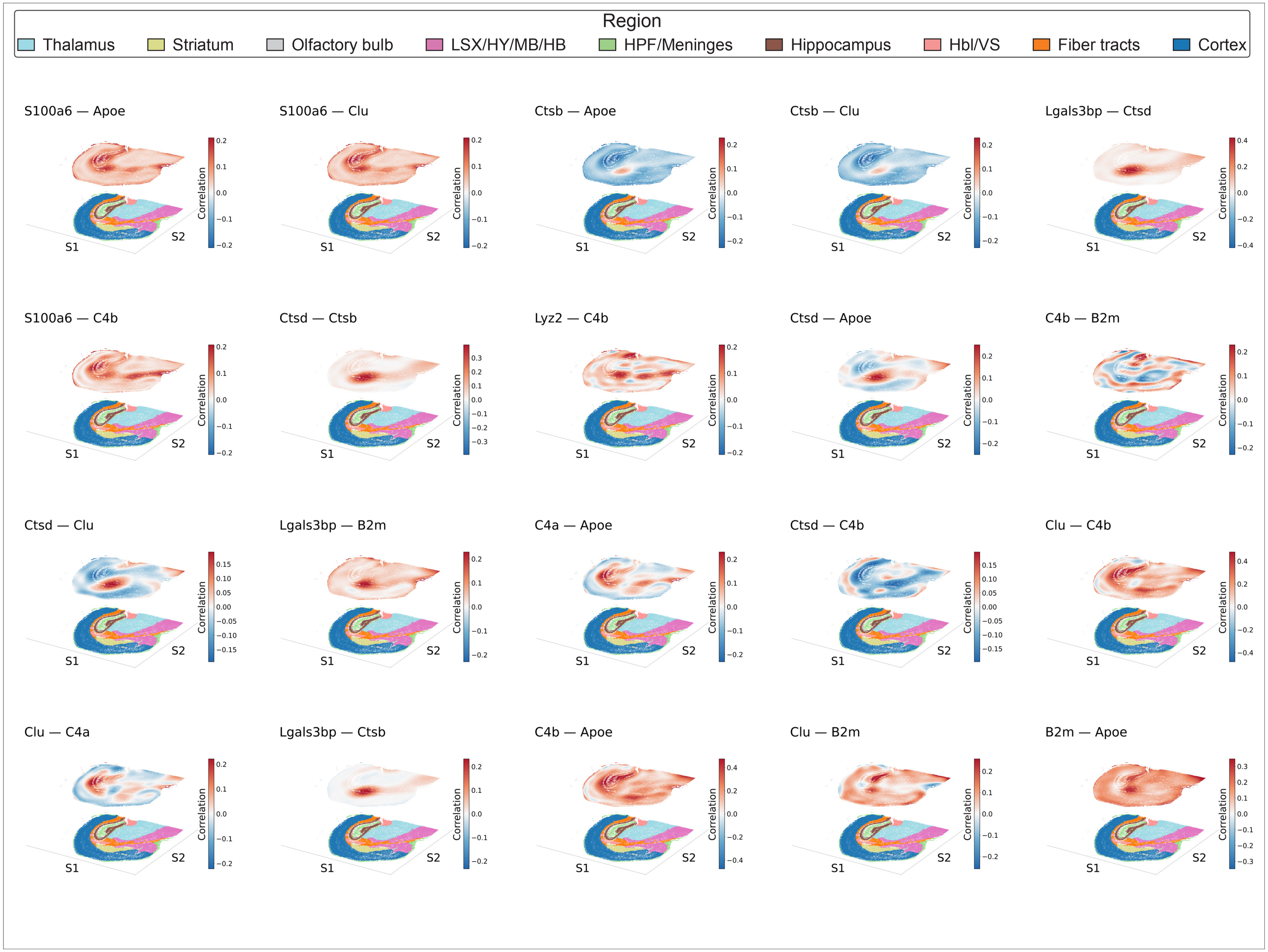

Figure S20: As in Figure S19, for the wild-type (WT) mouse brain. Panels show the estimated spatially varying correlation for the top 20 edges by sARI.

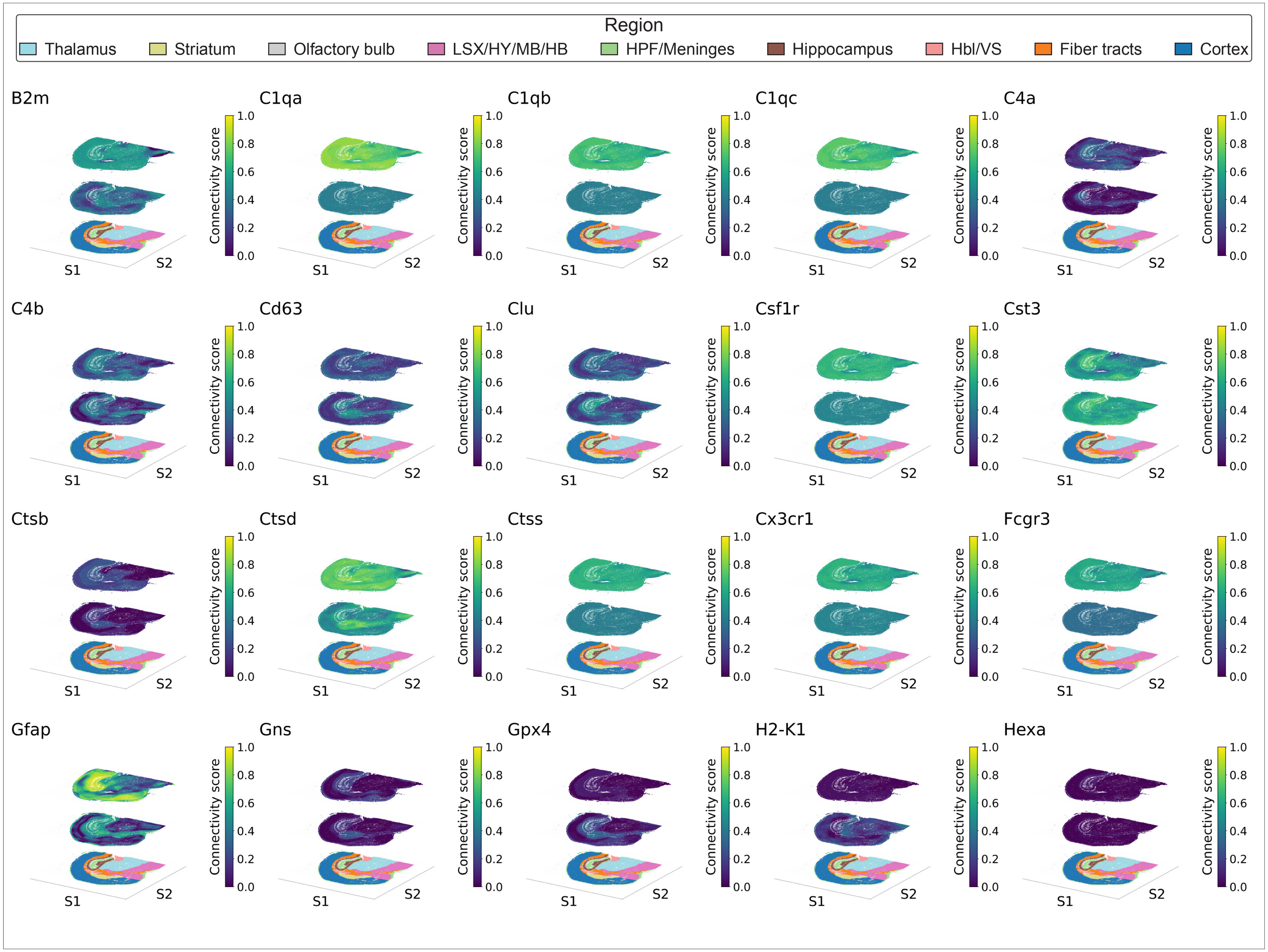

Figure S21: Gene-level spatially varying connectivity comparison between MUT and WT mouse brains. Each panel maps the connectivity score of a single gene across the tissue samples, summarizing how strongly that gene participates in the local co-expression network, with higher scores in yellow/green and lower scores in purple.

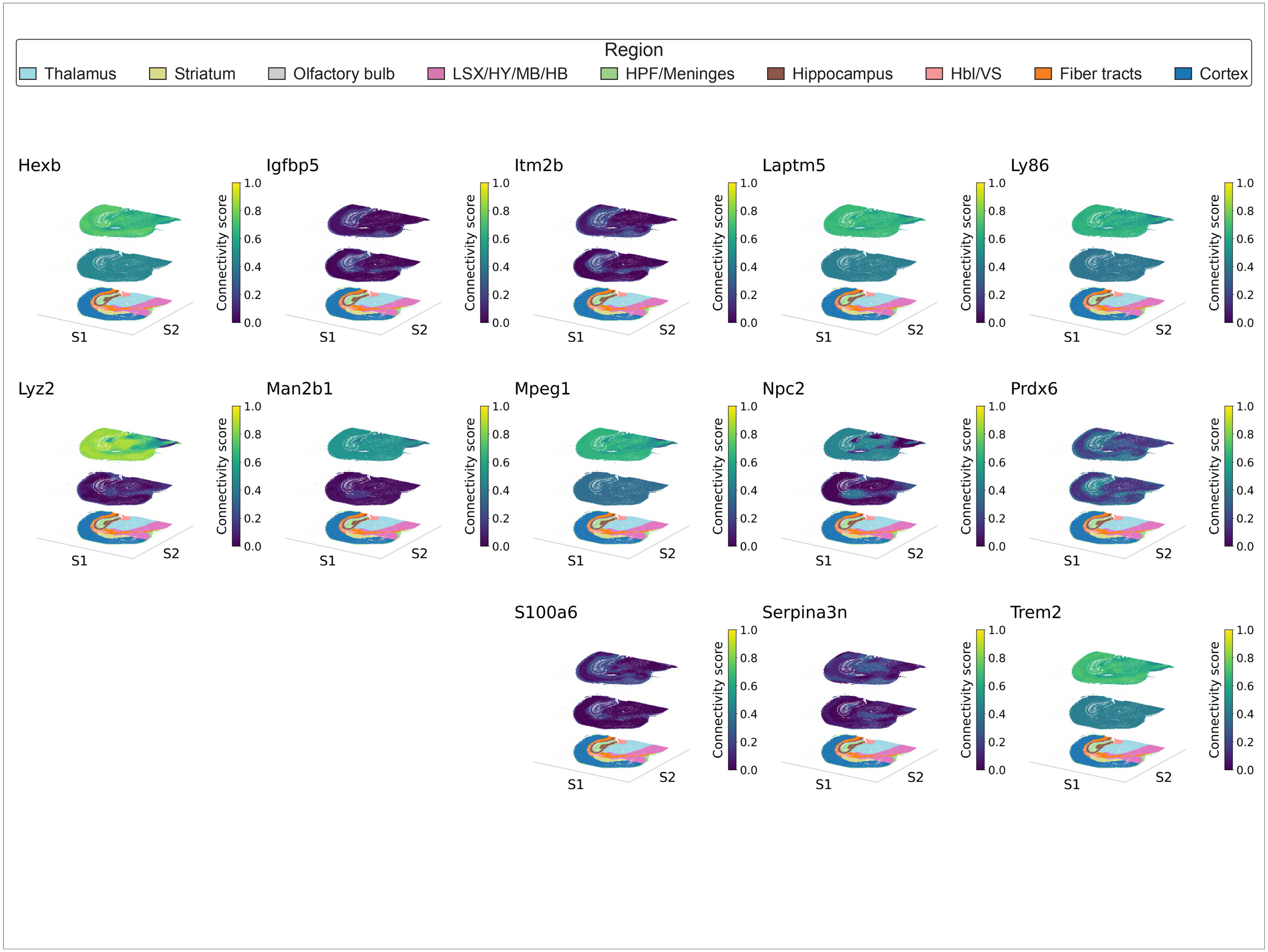

Figure S22: As in Figure S21, for the remaining genes in the pathway; each panel shows the spatially varying connectivity score of a single gene, allowing comparison of gene-level connectivity between genotypes.

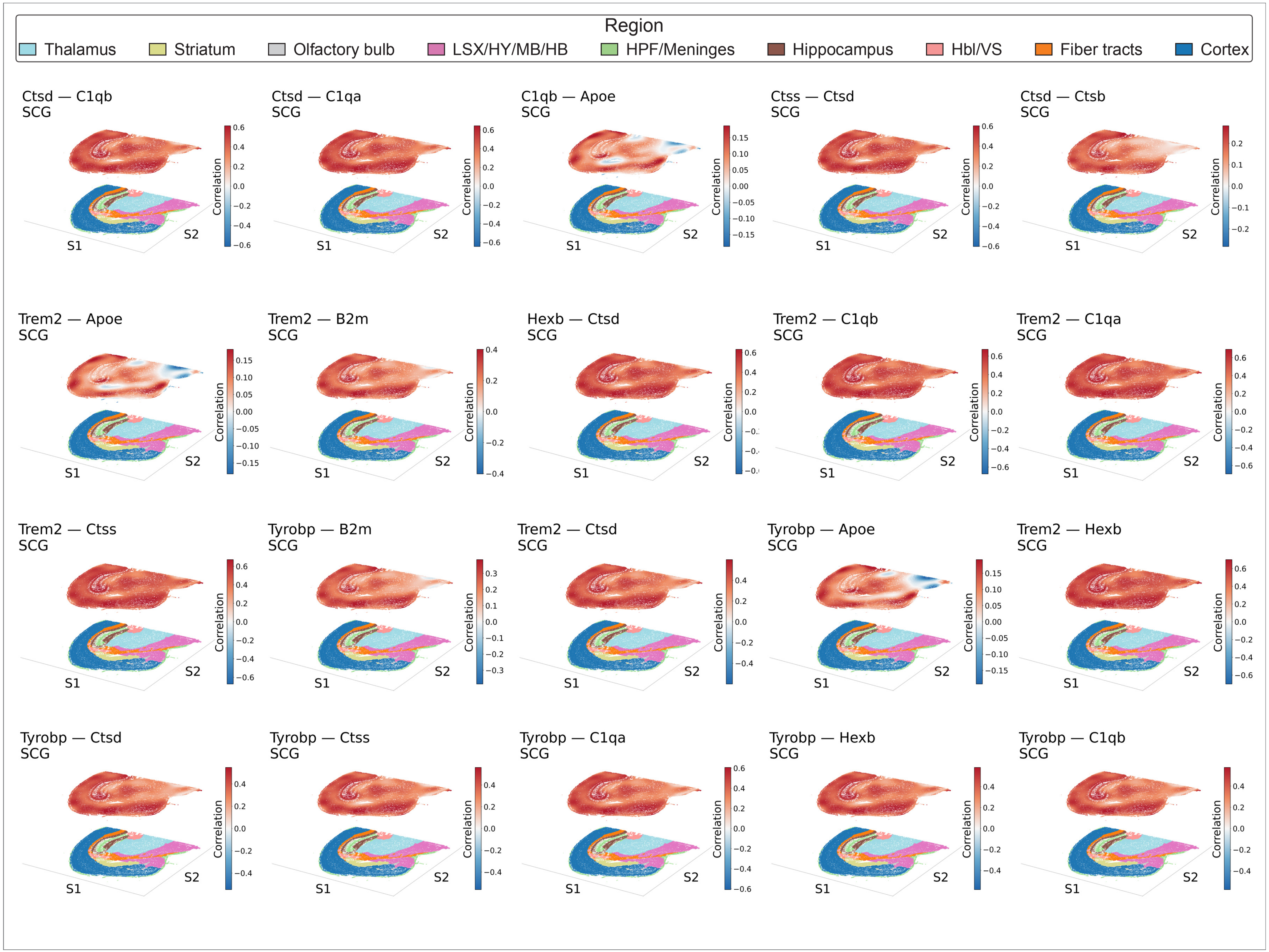

Figure S23: Spatially varying co-expression edges for selected gene pairs in the Alzheimer's disease mouse-model brain (mutant, MUT). Each panel maps the estimated posterior mean correlation of one gene pair across a coronal section, with red and blue denoting positive and negative co-expression. Anatomical regions are indicated in the legend. Edges displayed are among gene pairs identified as being co-expressed in the literatureChen et al. (2020).

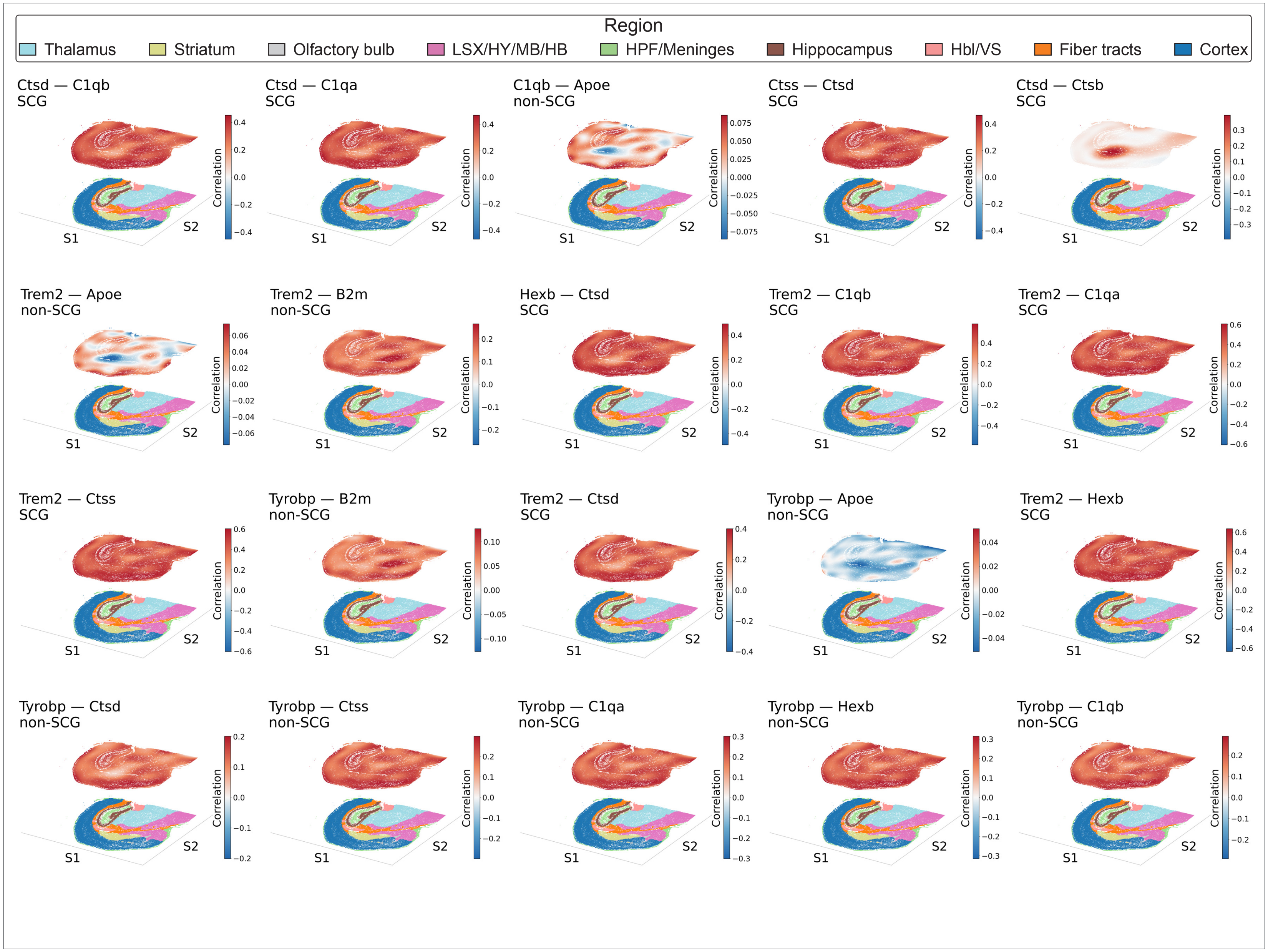

Figure S24: Figure S23, for the wild-type (WT) mouse brain. Panels show the estimated spatially varying correlation for the gene pairs identified as being co-expressed in the literatureChen et al. (2020)
